# SARS-CoV-2 mechanistic correlates of protection: insight from modelling response to vaccines

**DOI:** 10.1101/2021.10.29.466418

**Authors:** Marie Alexandre, Romain Marlin, Mélanie Prague, Séverin Coleon, Nidhal Kahlaoui, Sylvain Cardinaud, Thibaut Naninck, Benoit Delache, Mathieu Surenaud, Mathilde Galhaut, Nathalie Dereuddre-Bosquet, Mariangela Cavarelli, Pauline Maisonnasse, Mireille Centlivre, Christine Lacabaratz, Aurelie Wiedemann, Sandra Zurawski, Gerard Zurawski, Olivier Schwartz, Rogier W Sanders, Roger Le Grand, Yves Levy, Rodolphe Thiébaut

## Abstract

The definition of correlates of protection is critical for the development of next generation SARS-CoV-2 vaccine platforms. Here, we propose a new framework for identifying mechanistic correlates of protection based on mathematical modelling of viral dynamics and data mining of immunological markers. The application to three different studies in non-human primates evaluating SARS-CoV-2 vaccines based on CD40-targeting, two-component spike nanoparticle and mRNA 1273 identifies and quantifies two main mechanisms that are a decrease of rate of cell infection and an increase in clearance of infected cells. Inhibition of RBD binding to ACE2 appears to be a robust mechanistic correlate of protection across the three vaccine platforms although not capturing the whole biological vaccine effect. The model shows that RBD/ACE2 binding inhibition represents a strong mechanism of protection which required significant reduction in blocking potency to effectively compromise the control of viral replication.

**One Sentence Summary:** A framework for modelling the immune control of viral dynamics is applied to quantify the effect of several SARS-CoV-2 vaccine platforms and to define mechanistic correlates of protection.

## INTRODUCTION

There is an unprecedented effort for SARS-CoV-2 vaccine development with 294 candidates currently evaluated *(1)*. However, variants of concern have emerged before the vaccine coverage was large enough to control the pandemics *(2)*. Despite a high rate of vaccine protection, these variants might compromise the efficacy of current vaccines *(3–6)*. Control of the epidemic by mass vaccination may also be compromised by unknown factors such as long-term protection and the need of booster injections in fragile, immuno-compromised, elderly populations, or even for any individual if protective antibody levels wane. Furthermore, the repeated use of some of the currently approved vaccine could be compromised by potential adverse events or by immunity against vaccine viral vectors *(7)*. Finally, the necessity to produce the billions of doses required to vaccinate the world’s population also explains the need to develop additional vaccine candidates.

The identification of correlates of protection (CoP) is essential to accelerate the development of new vaccines and vaccination strategies *(8, 9)*. Binding antibodies to SARS-CoV-2 and *in vitro* neutralization of virus infection are clearly associated with protection *(10–13)*. However, the respective contribution to virus control *in vivo* remains unclear *(14)*, and many other immunological mechanisms may also be involved, including other antibody-mediated functions (antibody-dependent cellular cytotoxicity, antibody-dependent complement deposition, antibody-dependent cellular phagocytosis *(11, 15, 16)*), as well as T cell immunity *(17)*. Furthermore, correlates of protection may vary between the vaccine platforms *(18–21)*.

Non-human primate (NHP) studies offer a unique opportunity to evaluate early markers of protective response *(22, 23)*. Challenge studies in NHP allow the evaluation of vaccine impact on the viral dynamics in different tissue compartments (upper and lower respiratory tract) from for day one to virus exposure *(11, 15, 24)*. Such approaches in animal models may thus help to infer, for example, the relation between early viral events and disease or the capacity to control secondary transmissions.

Here, we propose a novel model-based framework to evaluate i) the immune mechanism involved in the vaccine response, and ii) the markers capturing this/these effect(s) leading to identification of mechanisms of protection and definition of mechanistic CoP *(25)*. First, we present a mechanistic approach based on ordinary differential equation (ODE) models reflecting the virus-host interaction *(26–29)*. The proposed model includes several new aspects refining the modeling of viral dynamics *in vivo*, in addition to the integration of vaccine effect. A specific inoculum compartment allows distinguishing the virus coming from the challenge inoculum and the virus produced *de novo*, which is a key point in the context of efficacy provided by antigen specific pre-existing immune effectors induced by the vaccine. Then, an original data mining approach is implemented to identify the immunological biomarkers associated with specific mechanisms of vaccine-induced protection.

We apply our approach to a recently published study *(30)* testing a protein-based vaccine targeting the receptor-binding domain (RBD) of the SARS-CoV-2 spike protein to CD40 (αCD40.RBD vaccine). Targeting vaccine antigens to Dendritic Cells via the surface receptor CD40 represents an appealing strategy to improve subunit-vaccine efficacy *(31–34)* and for boosting natural immunity in SARS-CoV-2 convalescent NHP.

We show that immunity induced by natural SARS-CoV-2 infection, as well as vaccine-elicited immune responses contribute to viral load control by i) blocking new infection of target cells and ii) by increasing the loss of infected cells. The modelling showed that antibodies inhibiting binding of RBD domain to ACE2 correlated with blockade of new infections and RBD binding antibodies correlate with the loss of infected cells, reflecting importance of additional antibody functionalities. The role of RBD/ACE2 binding inhibition has been confirmed in two other vaccine platforms.

## RESULTS

### A new mechanistic model fits the *in vivo* viral load dynamics in nasopharyngeal and tracheal compartments

The mechanistic model aims at capturing the viral dynamics following challenge with SARS-CoV-2 virus in NHP. For that purpose, we used data obtained from 18 cynomolgus macaques involved in the vaccine study reported by Marlin et al *(30)* and exposed to a high dose (1×10^6^ pfu) of SARS-CoV-2 administered via the combined intra-nasal and intra-tracheal route. The viral dynamics during the primary infections were characterized by a peak of genomic RNA (gRNA) production three days after infection, followed by a decrease toward undetectable levels beyond day 15 **(Figure S1)**. At the convalescent phase (median 24 weeks after the primary infection), 12 macaques were challenged with SARS-CoV-2 a second time, four weeks after being randomly selected to receive either a placebo (n=6) or a single injection of the αCD40.RBD vaccine (n=6) **(Figure 1A)**. A third group of 6 naïve animals were infected at the same time. Compared to this naïve group, viral dynamics were blunted following the second challenge of convalescent animals with the lowest viral load observed in vaccinated animals **(Figure 1B, S2)**.

**Fig. 1.**
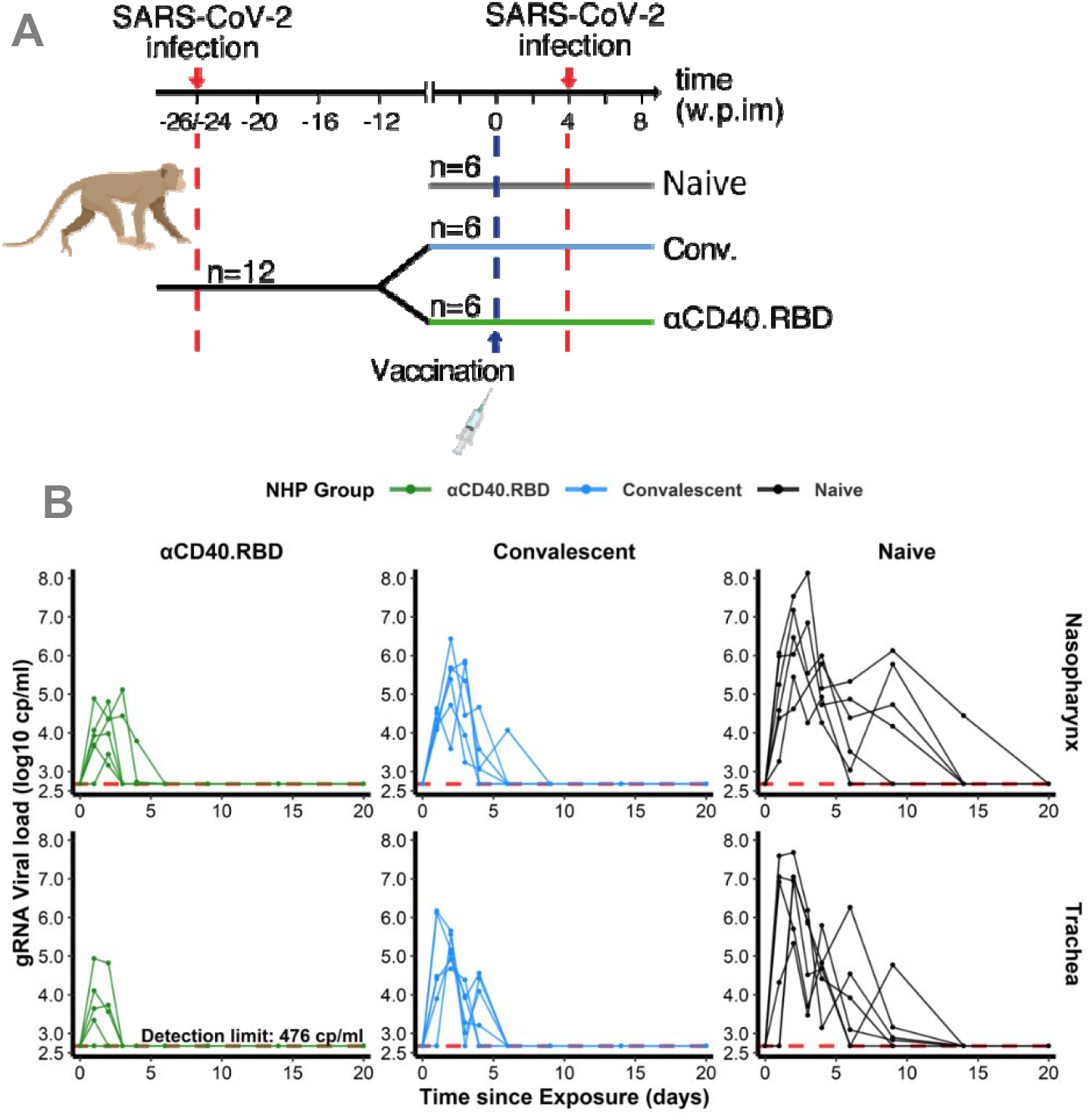
Design of the study 1 and viral dynamics. **(A)** *Study design*. Cynomolgus macaques (*Macaca fascicularis*), aged 37-58 months (8 females and 13 males). 24-26 weeks post infection with SARS-CoV-2, twelve of these animals were randomly assigned in two experimental groups. The convalescent vaccinated group (n=6) received 200 μg of αCD40.RBD vaccine. The other six convalescent animals were used as controls. Additional six age matched (43.7 months +/-6.76) cynomolgus macaques from same origin were included in the study as controls naive from any exposure to SARS-CoV-2. Four weeks after immunization, all animals were exposed to a total dose of 10^6^ pfu of SARS-CoV-2 virus via the combination of intra-nasal and intra-tracheal routes. **(B)** Individual log10 transformed gRNA viral load dynamics in nasopharyngeal swabs (top) and tracheal swabs (bottom) after the initial exposure to SARS-CoV-2 in naive macaques (black, right) and after the second exposure in convalescent (blue, middle) and αCD40.RBD-vaccinated convalescent (green, left) groups. Horizontal red dashed lines indicate the limit of quantification.

We developed a mathematical model to better characterize the impact of the immune response on the viral gRNA and subgenomic RNA (sgRNA) dynamics, adapted from previously published work *(26, 27, 35)*, which includes uninfected target cells (T) that can be infected (I_1_) and produce virus after an eclipse phase (I_2_). The virus generated can be infectious (V_i_) or non-infectious (V_ni_). We completed the model by a compartment for the inoculum to distinguish between the injected virus (V_s_) and the virus produced *de novo* by the host (V_i_ and V_ni_). The viral dynamics in the two compartments, the nasopharynx and the trachea, were jointly considered **(Figure 2A)**. Using the gRNA and sgRNA viral loads, we estimated the viral infectivity (β), the viral production rate (p) and the loss rate of infected cells (δ). We assumed that gRNA and sgRNA were proportional to the free virus and the infected cells, respectively. The duration of the eclipse phase, the clearance of the free virus from the inoculum and produced *de novo* were estimated separately by profile likelihood. The infectivity rate (0.95×10^−6^ (copies/ml)^-1^ day^-1^), the loss rate of infected cells (1.04 day^-1^), the eclipse phase (3 day^-1^) estimations in naïve animals were in the range of previously reported modelling results *(26, 27)*. Here, we distinguished the clearance of the inoculum which was much higher (20 virions day^-1^) as compared to the clearance of the virus produced *de novo* (3 virions day^-1^). Furthermore, the viral production by each infected cells was estimated to be higher in the nasopharyngeal compartment (12.1 10^3^ virions/cell/day) as compared to the tracheal compartment (0.92 10^3^ virions/cell/day). These estimations are in agreement with the observation of the intense production of viral particles by primary human bronchial epithelial cells in culture *(36)*. By allowing parameters to differ between animals (through random effects), the variation of cell infectivity and of the loss rate of infected cells captured the observed variation of the dynamics of viral load. The variation of those parameters could be partly explained by the group to which the animals belong reducing the unexplained variability of the cell infectivity by 66% and of the loss rate of infected cells by 54% **(Table S1)**. The model fitted well the observed dynamics of gRNA and sgRNA **(Figure 2B)**.

**Fig. 2.**
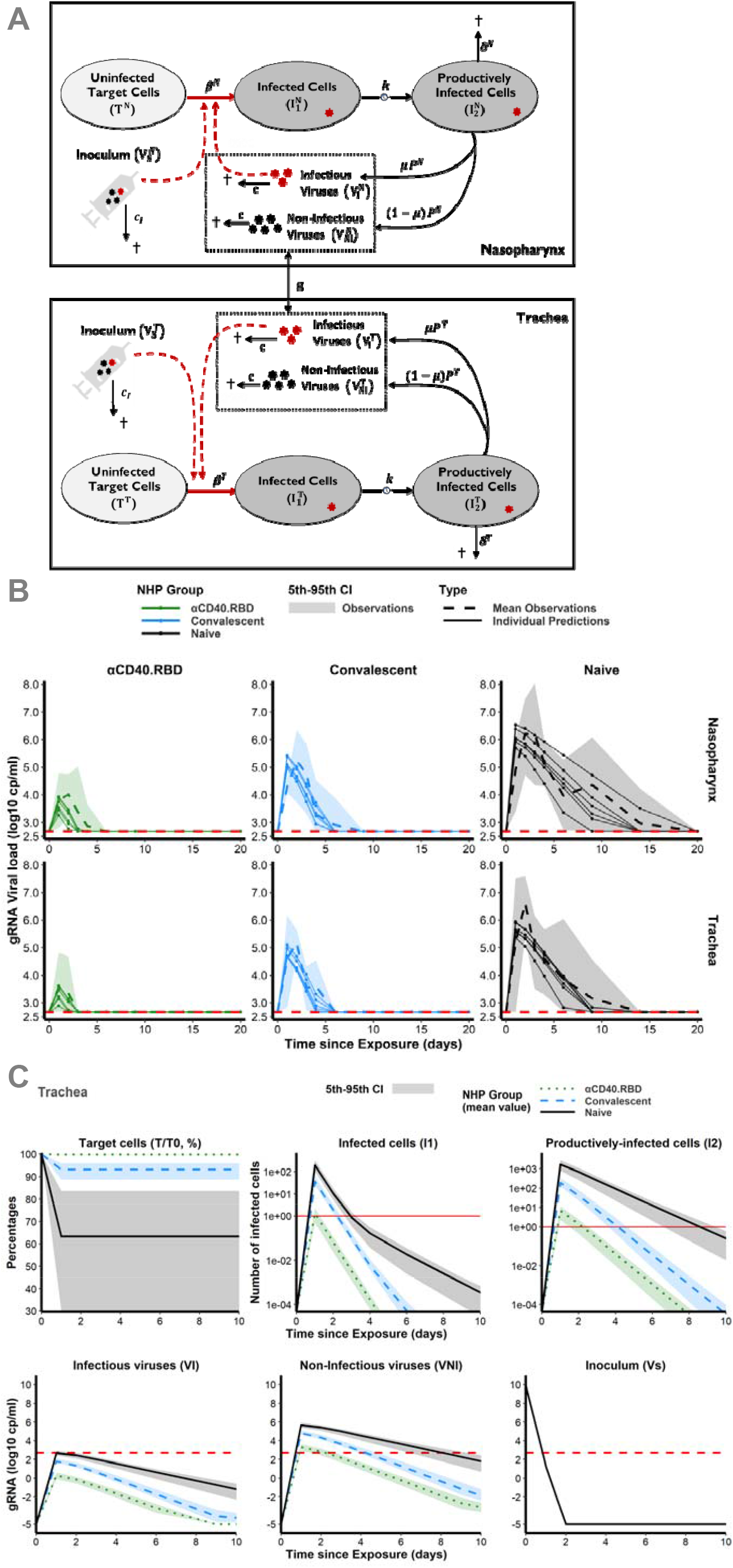
Mechanistic modelling. **(A)** Description of the model in the two compartments: the nasopharynx and the trachea. **(B)** Model fit to the log10 transformed observed gRNA viral loads in nasopharyngeal (top) and tracheal (bottom) compartments after the initial exposure to SARS-CoV-2 in naive macaques (black, right) and after the second exposure in convalescent (blue, middle) and vaccinated (green, left) animals. Solid thin lines indicate individual dynamics predicted by the model adjusted on the effect of group. Thick dashed lines indicate mean viral loads over time. Shaded areas indicate the 95% confidence interval. Horizontal red dashed lines indicate the limit of quantification. **(C)** Model predictions of unobserved quantities in the tracheal compartment for naive (black, solid lines), convalescent (blue, dashed lines) and vaccinated (green, dotted lines) animals: target cells as percentage of the value at the challenge (top, left), infected cells (top, middle), productively infected cells (top, right), inoculum (bottom, right), infectious (bottom, left) and non-infectious virus (bottom, middle). Thick lines indicate mean values over time within each group. Shaded areas indicate the 95% confidence interval. Horizontal dashed red lines indicate the limit of quantification and horizontal solid red lines highlight the threshold of one infected cell.

### Modelling of the dynamics of viral replication argues for the capacity of αCD40.RBD vaccine to block virus entry into host cells and to promote the destruction of infected cells

We distinguish the respective contribution of the vaccine effect and post-infection immunity on the reduction of the cell infection rate and the increase of the clearance of infected cells. Because blocking *de novo* infection and promoting the destruction of infected cells would lead to different viral dynamics profile **(Figure S3)**, we were able to identify the contribution of each mechanism by estimating the influence of the vaccine compared to placebo or naive animals on each model parameter. The αCD40.RBD vaccine reduced by 99.6% the infection of target cells in the trachea compared to the naïve group. The estimated clearance of infected cells was 1.04 day^-1^ (95% CI 0.75; 1.45) in naïve macaques. It was increased by 80% (1.86/day^-1^) in the convalescent macaques vaccinated by αCD40.RBD or not.

The mechanistic model allows predicting the dynamics of unobserved compartments. Hence, a very early decrease of the target cells (all cells expressing ACE2) as well as of the viral inoculum which fully disappeared from day 2 onward were predicted **(Figure 2C)**. In the three groups, the number of infected cells as well as infectious viral particles increased up to day 2 and then decreased. We show that this viral dynamic was blunted in the vaccinated animals leading to a predicted maximum number of infectious viral particles in the nasopharynx and the trachea below the detection threshold **(Figure 2C)**. The number of target cell levels would be decreased by the infection in the naïve and the convalescent groups, whereas it would be preserved in vaccinated animals.

### The RBD-ACE2 binding inhibition is the main mechanistic CoP explaining the effect of the αCD40.RBD vaccine on new cell infection

In our study *(30)*, an extensive evaluation of the immunological response has been performed with quantification of spike binding antibodies, antibodies inhibiting the attachment of RBD to ACE2, antibodies neutralizing infection, SARS-CoV-2-specific CD4^+^ and CD8^+^ T cells producing cytokines and serum cytokine levels **(Figure 3, S4, S5, S6)**. Therefore, based on our mechanistic model we investigated if any of these markers could serve as a mechanistic CoP. Such a CoP should be able to capture the effect of the natural immunity following infection, associated or not to the vaccine (group effect) estimated on both the rate of cell infection and the rate of the loss of infected cells. To this aim, we performed a systematic screening by adjusting the model for each marker and we compared these new models to the reference model adjusted for the groups (See supplementary information for a detailed description of the algorithm). We demonstrate that the RBD-ACE2 binding inhibition measure is sufficient to capture most of the effect of the groups on the infection of target cells **(Figure 4A, 4B)**. The integration of this marker in the model explains the variability of the cell infection rate with greater certainty than the group of intervention, reducing the unexplained variability by 87% compared to 66% **(Table S1)**. The marker actually takes into account the variation between animals within the same group. Hence, it suggests that the levels of anti-RBD antibodies induced by the vaccine that block attachment to ACE2 are highly efficient at reflecting the neutralization of new infections *in vivo*. Furthermore, when taking into account the information provided by the RBD-ACE2 binding inhibition assay, the effect of the group of intervention was no longer significant **(Table S1)**. Finally, we looked at the estimated infection rate according to the inhibition binding assay in every animal **(Figure 4C)**. The values were not overlapping at all, distinguishing clearly the vaccinated and unvaccinated animals.

**Fig. 3.**
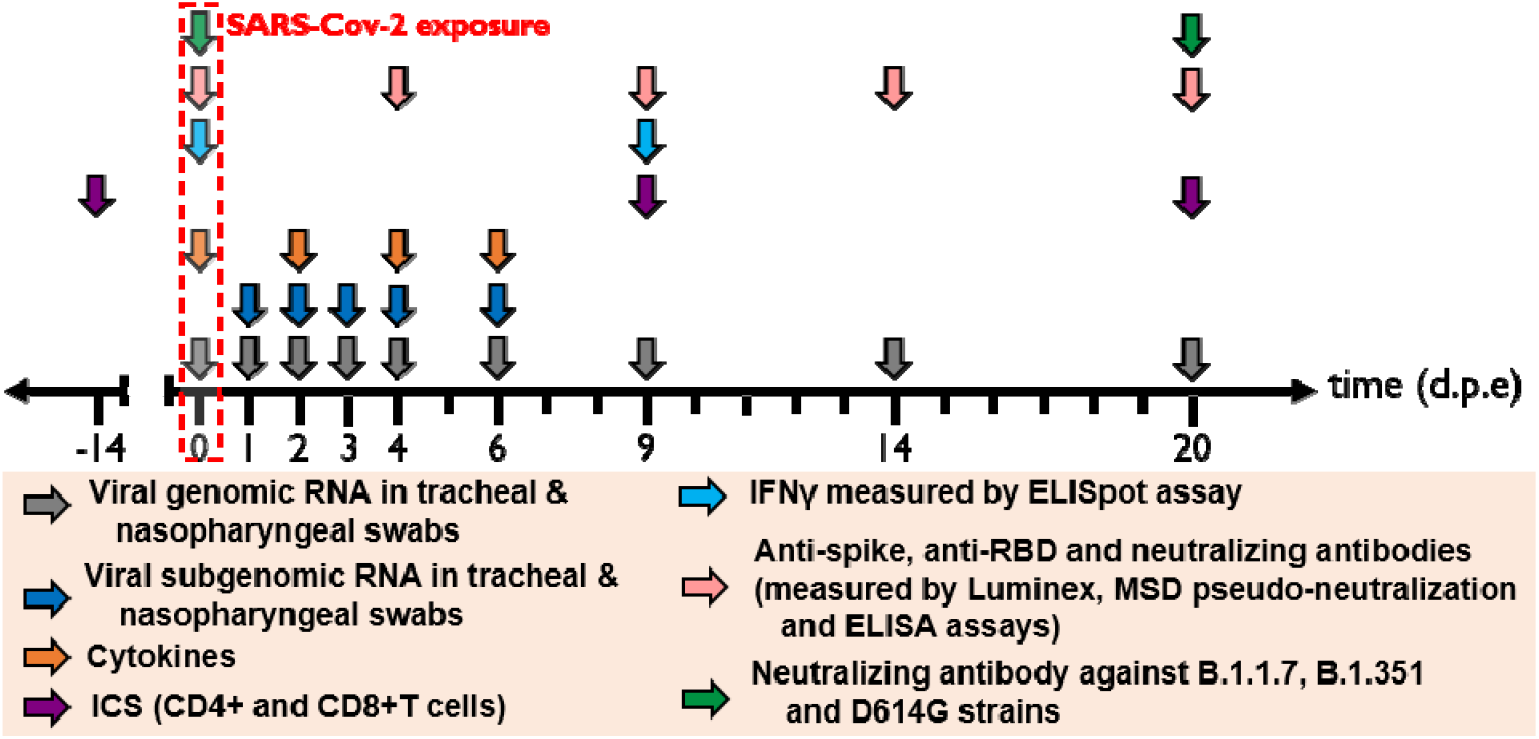
Harvest times and measurements. Nasopharyngeal and tracheal fluids, were collected at 0, 1, 2, 3, 4, 6, 9, 14 and 20 days post exposure (d.p.e) while blood was taken at 0, 2, 4, 6, 9, 14 and 20 d.p.e. Genomic and subgenomic viral loads were measured by RT-qPCR. Anti-Spike IgG sera were titrated by multiplex bead assay, Anti-RBD and anti-Nucleocapside (N) IgG were titrated using a commercially availabl multiplexed immunoassay developed by Mesoscale Discovery (MSD, Rockville, MD). The MSD pseudo-neutralization assay was used to measure antibodies neutralizing the binding of the spike protein and RBD to the ACE2 receptor. Neutralizing antibodies against B.1.1.7, B.1.351 and D614G strains were measured by S-Fuse neutralization assay and expressed as ED50 (Effective dose 50%). T-cell responses were characterized as the frequency of PBMC expressing cytokines (IL-2, IL-17 a, IFN-_γ_, TNF-a, IL-13, CD137 and CD154) after stimulation with S or N sequence overlapping peptide pools. IFN-_γ_ ELISpot assay of PBMCs were performed on PBMC stimulated with RBD or N sequence overlapping peptide pools and expressed as spot forming cell (SFC) per 1.0×10^6^ PBMC.

**Fig. 4.**
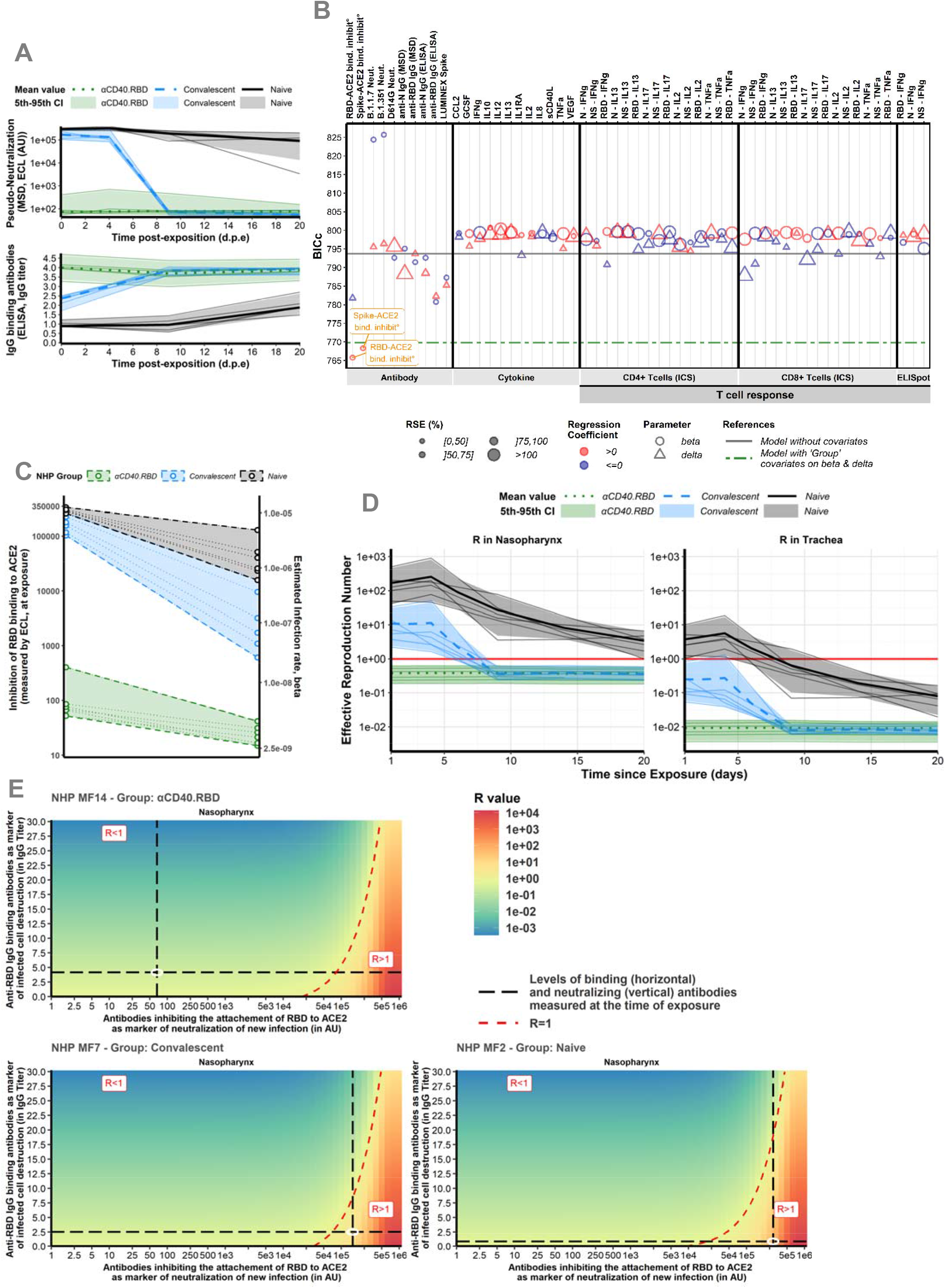
Immune markers. **(A)** *Dynamics of biomarker selected as mCoP*. Quantification of antibodies inhibiting RBD-ACE2 binding, measured by the MSD pseudo-neutralization assay (ECL, in AU) (top) and anti-RBD IgG titrated by ELISA assay (in IgG titer) (bottom). Thin lines represent individual values. Thick lines indicate medians within naïve (black, solid line), convalescent (blue, dashed line) and αCD40.RBD-vaccinated convalescent (green, dotted line) animals. Shaded areas indicate 5th-95th confidence intervals. **(B)** *Systematic screening of effect of the markers*. For every single marker, a model has been fitted to explore whether it explains the variation of the parameter of interest better or as well than the group indicator. Parameters of interest were β, the infection rate of ACE2+ target cells, and δ, the loss rate of infected cells. Models were compared according to the Bayesian Information Criterion (BIC), the lower being the better. The green line represents the reference model that includes the group effect (naive/convalescent/vaccinated) without any adjustment for immunological marker (see **Figure 3** for more details about measurement of immunological markers). **(C)** *Thresholds of inhibition of RBD-ACE2 binding*. Estimated infection rate (in (copies/mL)^-1^ day^-1^) of target cells according to the quantification of antibodies inhibiting RBD-ACE2 (in ECL) at exposure. Thin dotted lines and circles represent individual values of infection rates (right axis) and neutralizing antibodies (left axis). Shaded areas delimit the pseudo-neutralization / viral infectivity relationships within each group. **(D)** *Reproduction rate over time*. Model predictions of the reproduction rate over time in the trachea (right) and nasopharynx (left). The reproduction rate is representing the number of infected cells from one infected cell if target cells are unlimited. Below one, the effective reproduction rate indicates that the infection is going to be cured. Horizontal solid red lines highlight the threshold of one. Same legend than A). **(E)** *Conditions for controlling the infection*. Basic reproduction rate at the time of the challenge according to the levels of antibodies inhibiting RBD-ACE2 binding (the lower the better) and of anti-RBD IgG binding antibodies (the higher the better) assuming they are mechanistic correlates of blocking new cell infection and promoting infected cell death, respectively. The red area with R>1 describes a situation where the infection is spreading. The green area with R<1 describes a situation where the infection is controlled. The dotted red line delimitates the two areas. Black long dashed lines represent the values of neutralizing and binding antibodies measured at exposure. Observed values for three different animals belonging to the naive (bottom, right), convalescent (bottom, left) and vaccinated (top, left) groups are represented.

In the next step, several markers (IgG binding anti-RBD antibodies, CD8^+^ T cells producing IFN-γ) appeared to be associated to the rate of loss of infected cells (**Figure S7A**). Both specific antibodies and specific CD8^+^ T cells are mechanisms commonly considered important for killing infected cells. We retained the anti-RBD binding IgG Ab that were positively associated to the increase of the loss of infected cells. For unknown reason the IFN-γ response was high in unstimulated conditions in the naïve group. Thus, although this marker was associated with a decrease of the loss rate of infected cells, it appears essentially here as an indicator of the animal group. Further studies would be needed to fully confirm the place of IFN-γ response as a mechanistic marker.

A large part of the variation of the infection rate (71%) and loss rate of infected cells (60%) were captured by the two markers of CoP: the RBD-ACE2 binding inhibition and the anti-RBD binding Ab concentration. Using the estimated parameters, the effective reproduction rate could be calculated (R) which is representing the number of cells secondarily infected by virus from one infected cell **(Figure 4D)**. When looking at this effective reproduction rate according to the groups, the vaccinated animal presented from the first day of challenge an effective R below 1 meaning that no propagation of the infection started within the host. These results were consistent when taking the value of RBD-ACE2 binding inhibition at the time of the challenge without considering the evolution of the inhibition capacity over time **(Figure S7B)**. This means that the dynamics of the viral replication is impacted very early during the infection process in immunized animals and that vaccinated animals were protected from the beginning by the humoral response. Then, we looked at the threshold of the markers of interest leading to the control of the within-host infection (as defined by R<1) which was around 30 000 AU for the RBD-ACE2 binding inhibition assay. For the animals in the naive and the convalescent groups, the observed values of binding inhibition measured by ECL RBD (the lower the better) and of IgG anti-RBD binding antibodies (the higher the better) led to R>1, whereas in vaccinated animals, the value of ECL RBD led to R<1. Therefore, our modeling study shows that the inhibition of binding of RBD to ACE2 by antibodies is sufficient to control initial infection of the host (**Figure 4E**). According to the observed value of ECL RBD in vaccinated animals (e.g., 66 AU in **Figure 4E**), a decrease of more than 2 log10 of the inhibition capacity (to reach 81 000 AU), due to variant of concern (VoC) or waning of immunity, would have been necessary to impair the control of the within-host infection. Moreover, a decrease of the neutralizing activity (i.e., increased ECL) could be compensated by an increase of cell death as measured by an increase of binding IgG anti-RBD as a surrogate. As an example, increasing IgG anti-RBD from 2.5 to 10 in the animal MF7 of the convalescent group would lead to a control of the infection. In conclusion, the αCD40.RBD vaccine-elicited humoral response leads to the blockade of new cell infection that is well captured by measure of the inhibition of attachment of the virus to ACE2 through the RBD domain of the spike protein. Hence, the inhibition of binding of RBD to ACE2 is a promising mechanistic CoP. Indeed, this CoP fulfils the three criteria of leading to the best fit (lower BIC), the best explanation of inter-individual variability, and fully captured the effect of the group of intervention.

### The model revealed the same CoP related to another protein-based vaccine but not with mRNA-1273 vaccine

We took the opportunity of another study testing a two-component spike nanoparticle vaccine performed in the same laboratory and using the same immune and virological assays *(37)* for applying the proposed model and methodology. In this study, 6 animals were vaccinated and compared to 4 naive animals (**Figure S8A, S8B**). The good fit of the data (**Figure S8C, S8D**) allows for estimating the effect of the vaccine that appeared here also to decrease the transmission rate (by 99%) and increase the clearance of the infected cells by 79%. Looking at the best mechanistic CoP following the previously described strategy, we ended here again with the inhibition of RBD binding to ACE2 as measured by ECL RBD. In fact, this marker measured at baseline before challenge responded to the three criteria: i) it led to the best model in front of a model adjusted for group effect, ii) it rendered the group effect non-significant and iii) it explained around 71% of the transmission rate variability, compared to 65% of variability explained by the groups. Interestingly, here again, the inhibition assay led to a clear separation of the estimated rate of transmission between vaccinees and the placebo group (**Figure S8E**).

Finally, we applied our approach to a published NHP study performed to evaluate several doses of mRNA-1273 vaccine *(24)*. Using available data, we compared the viral dynamics in the 100 μg, 10 μg and placebo group. We started from the same model as defined previously. We estimated a reduction of transmission rate by 97% but we did not find any additional effect. Looking at potential mechanistic CoP, we retained neutralization as measured on live cells with Luciferase marker. Although this marker led to the best fit and replaced the group effect (which was non-significant after adjustment for the marker), it explained only 15% of the variability of estimated transmission rate, while 19% were explained by the groups.

In conclusion, we demonstrated, based upon challenge studies in NHP vaccinated with two different protein-based vaccine platforms that both vaccines lead to the blockade of new cell infection. Neutralizing antibodies likely represent a consistent mechanistic correlate of protection. This could change across vaccine platforms especially because mechanisms of action are different.

## DISCUSSION

We propose a novel framework to explore the mechanistic effects of vaccines and to assess the quality of markers as mechanistic CoP (mCoP) that we applied to SARS-CoV-2 vaccines. This model showed that neutralizing and binding antibodies, elicited by a non-adjuvanted protein-based vaccine targeting the RBD of spike to the CD40 receptor of antigen presenting cells are reliable mCoP. Interestingly, we found the simpler and easier to standardize and realize binding inhibition assay may be more relevant to use as a correlate of protection than cell-culture neutralization assays. This result has been replicated in another study testing a nanoparticle spike vaccine. The model was able to capture the effect of the vaccines on the reduction of the rate of infection of target cells and identified additional effects of vaccines beyond neutralizing antibodies. This latter consisted of increasing the loss rate of infected cells which was better reflected by the IgG binding antibodies and CD8^+^ T cell responses in the case of the CD40-targeting vaccine. One limitation of our study is that the prediction potential of our model relies on the range of the immune markers measured. However, our approach would allow a full exploitation of the data generated as in systems serology where non-neutralizing Ab functions, such as Ab-dependent cellular cytotoxicity (ADCC), Ab-dependent cellular phagocytosis (ADCP), Ab-dependent complement deposition (ADCD), and Ab-dependent respiratory burst (ADRB) are explored *(38)*. The role of ADCC in natural infection has been previously shown *(39)*, ADCD in DNA vaccine recipients *(11)* and with Ad26 vaccine *(40)*. Here, we extended significantly these data by modelling the viral dynamic, showing that two other protein-based vaccines exert an additional effect on infected cell death which relied on the level of IgG anti-RBD binding antibodies especially for the CD40.RBD targeting vaccine. Measurements of other non-neutralizing Ab functions would probably also capture this additional effect.

The next question after determining which marker is a valid mCoP is to define the concentration that leads to protection, looking for a threshold effect that will help to define an objective *(10, 41)*. In the context of SARS-CoV-2 virus, several emerged variants are leading to a significant reduction of viral neutralization as measured by various approaches. However, a 20-fold reduction of viral neutralization might not translate in 20-fold reduction of vaccine efficacy *(42)*. First, there are many steps between viral neutralization and the reduction of transmission or the improvement of clinical symptoms. Second, the consequences of a reduction of viral neutralization could be alleviated by other immunological mechanisms not compromised by the variant. In the context of natural immunity, when the level of neutralizing antibodies was below a protective threshold, the cellular immune response appeared to be critical *(17, 43)*. We showed with our model that an improvement of infected cell destruction could help to control the within-host infection and is quantitatively feasible.

The control of viral replication is the key for reducing transmission *(44, 45)* as well as disease severity *(46–48)*. According to our non-linear model linking the neutralization to the viral replication, a decrease of 4 to 20 fold in neutralization as described for the variants of concern *(4, 6)* is not enough, especially in the context of the response to CD40.RBD targeting vaccine, to compromise the control of viral replication. This potential limited impact of variants on the host viral dynamics should be associated to a reduced transmission of escape variants in vaccinated population as compared to wild type virus in the unvaccinated population *(49)*. The results showing a conserved effectiveness of mRNA vaccines in humans infected by the alpha or beta variants *(50)*, although a decrease of neutralization has been reported *(4)*, are consistent with this hypothesis. However, this is highly dependent upon the mode of action of currently used vaccines and if there is no new VoC compromising the neutralization in a much higher scale than what has been described to date *(51, 52)*. This may globally have an impact on the global burden of the pandemic, since the occurrence of variants within host is probably a rare event *(53)* more likely occurring in specific conditions *(54)* and therefore the strongest selection for vaccine-escape mutants occurs by transmission *(49)*. In the case of delta variants, a marked decrease of neutralization has been described *(55)* but the impact on vaccine effectiveness is less clear *(56)*.

The analysis performed extended significantly the observation of associations between markers as previously reported for SARS-CoV-2 vaccine *(11)* and other vaccines *(57)* because it allows a more causal interpretation of the effect of immune markers. However, our modelling approach requires the *in vivo* identification of the biological parameters under specific experimentations. On the other hand, the estimation of parameters included in our model also provided information on some aspect of the virus pathophysiology. Notably, we found an increased capacity of virion production in nasopharynx compared to the trachea which could be explained by the difference in target cells according to the compartment *(58)*.

In conclusion, the framework presented here based on a mathematical model of viral dynamics should help in better evaluating the effect of vaccines and defining mechanistic CoP. The application to two new promising SARS-CoV-2 vaccines revealed a combination of effects with a blockade of new cell infections and the destruction of infected cells. For these two vaccines, the antibody inhibiting the attachment of RBD to ACE2, appeared to be a very good surrogate of the vaccine effect on the rate of infection of new cells and therefore could be used as a mechanistic CoP. This modelling framework participates to the improvement of the understanding of the immunological concepts by adding a quantitative evaluation of the contributions of different mechanisms of control of viral infection. In terms of acceleration of vaccine development, our results may help to develop vaccines for “hard-to-target pathogens”, or to predict their efficacy in aging and particular populations *(59)*. It should also help in choosing vaccine dose, for instance at early development *(60)* as well as deciding if and when boosting vaccination is needed in the face of waning protective antibody levels *(61, 62)*.

## MATERIALS AND METHODS

### Experimental model and subjects details

Cynomolgus macaques (Macaca fascicularis), aged 37-66 months (18 females and 13 males) and originating from Mauritian AAALAC certified breeding centers were used in this study. All animals were housed in IDMIT facilities (CEA, Fontenay-aux-roses), under BSL2 and BSL-3 containment when necessary (Animal facility authorization #D92-032-02, Préfecture des Hauts de Seine, France) and in compliance with European Directive 2010/63/EU, the French regulations and the Standards for Human Care and Use of Laboratory Animals, of the Office for Laboratory Animal Welfare (OLAW, assurance number #A5826-01, US). The protocols were approved by the institutional ethical committee “Comité d’Ethique en Expérimentation Animale du Commissariat à l’Energie Atomique et aux Energies Alternatives” (CEtEA #44) under statement number A20-011. The study was authorized by the “Research, Innovation and Education Ministry” under registration number APAFIS#24434-2020030216532863v1.

### Evaluation of anti-Spike, anti-RBD and neutralizing IgG antibodies

#### Anti-Spike IgG were titrated by multiplex bead assay

Briefly, Luminex beads were coupled to the Spike protein as previously described *(63)* and added to a Bio-Plex plate (BioRad). Beads were washed with PBS 0.05% tween using a magnetic plate washer (MAG2x program) and incubated for 1h with serial diluted individual serum. Beads were then washed and anti-NHP IgG-PE secondary antibody (Southern Biotech, clone SB108a) was added at a 1:500 dilution for 45 min at room temperature. After washing, beads were resuspended in a reading buffer 5 min under agitation (800 rpm) on the plate shaker then read directly on a Luminex Bioplex 200 plate reader (Biorad). Average MFI from the baseline samples were used as reference value for the negative control. Amount of anti-Spike IgG was reported as the MFI signal divided by the mean signal for the negative controls.

*Anti-RBD and anti-Nucleocapside (N) IgG* were titrated using a commercially available multiplexed immunoassay developed by Mesoscale Discovery (MSD, Rockville, MD) as previously described *(64)*. Briefly, antigens were spotted at 200−400 μg/mL in a proprietary buffer, washed, dried and packaged for further use (MSD^®^ Coronavirus Plate 2). Then, plates were blocked with MSD Blocker A following which reference standard, controls and samples diluted 1:500 and 1:5000 in diluent buffer were added. After incubation, detection antibody was added (MSD SULFO-TAGTM Anti-Human IgG Antibody) and then MSD GOLDTM Read Buffer B was added and plates read using a MESO QuickPlex SQ 120MM Reader. Results were expressed as arbitrary unit (AU)/mL.

*Anti-RBD and anti-N IgG* were titrated by ELISA. The Nucleocapsid and the Spike RBD domain (Genbank # NC_045512.2) were cloned and produced in *E. Coli* and CHO cells, respectively, as previously described *(31)*. Antigens were purified on C-tag column (Thermo Fisher) and quality-controlled by SDS-PAGE and for their level of endotoxin. Antigens were coated in a 96 wells plates Nunc-immuno Maxisorp (Thermo Fisher) at 1 μg/mL in carbonate buffer at 4°C overnight. Plates were washed in TBS tween 0.05% (Thermo Fisher) and blocked with PBS 3% BSA for 2 hours at room temperature. Samples were then added, in duplicate, in serial dilution for 1 hour at RT. Non-infected NHP sera were used as negative controls. After washing, anti-NHP IgG coupled with HRP (Thermo Fisher) was added at 1:20,000 for 45 min at RT. After washing, TMB substrate (Thermo Fisher) was added for 15 min at RT and the reaction was stopped with 1M sulfuric acid. Absorbance of each well was measured at 450 nm (reference 570 nm) using a Tristar2 reader (Berthold Technologies). The EC_50_ value of each sample was determined using GraphPad Prism 8 and antibody titer was calculated as log (1/EC_50_).

*The MSD pseudo-neutralization assay* was used to measure antibodies neutralizing the binding of the spike protein to the ACE2 receptor. Plates were blocked and washed as above, assay calibrator (COVID-19 neutralizing antibody; monoclonal antibody against S protein; 200 μg/mL), control sera and test sera samples diluted 1:10 and 1:100 in assay diluent were added to the plates. Following incubation of the plates, an 0.25 μg/mL solution of MSD SULFO-TAGTM conjugated ACE-2 was added after which plates were read as above. Electro-chemioluminescence (ECL) signal was recorded.

### Viral dynamics modelling

The mechanistic approach we developed to characterize the impact of the immune response on the viral gRNA and sgRNA dynamics relies on a mechanistic model divided in three layers: firstly, we used a mathematical model based on ordinary differential equations to describe the dynamics in the two compartments, the nasopharynx and the trachea. Then we used a statistical model to take into account both the inter-individual variability and the effects of covariates on parameters. Finally, we considered an observation model to describe the observed log_10_ viral loads in the two compartments.

For the mathematical model, we started from previously published models *(26, 27, 35)* where nasopharynx and trachea were described by target cell limited models and we completed the model by adding a compartment for the inoculum to be able to distinguish between the injected virus (V_s_) and the virus produced *de novo* (V_i_ and V_ni_). Consequently, for each of the two compartments, the model included uninfected target cells (T) that can be infected (I_1_) either by infectious viruses (V_i_) or inoculum (Vs) at an infectivity rate β. After an eclipse phase, infected cells become productively infected cells (I_2_) that can produce virions at rate *P* and be lost at a per capita rate δ. The virions generated can be infectious (V_i_) with proportion μ while the (1-μ) remaining virions are non-infectious (V_ni_). Finally, free *de novo* produced virions and free virions from inoculum are respectively cleared at a rate *c* and *c*_*i*_. The model can be written as the following set of differential equations, where the superscript X denotes the compartment of interest (N, nasopharynx or T, trachea):

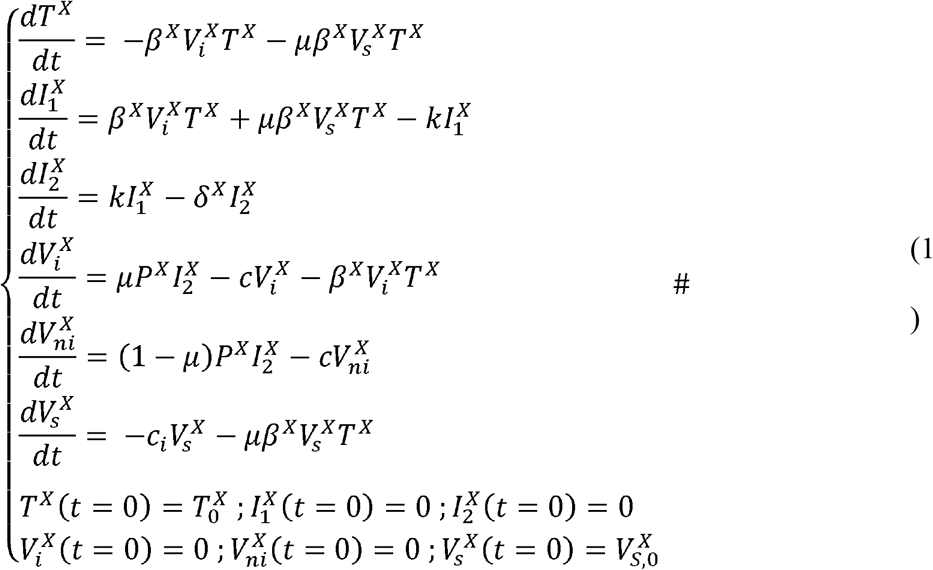

where *T*^*X*^(*t* = 0),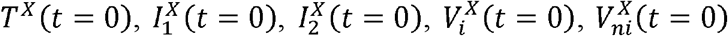 and 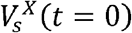 are the initial conditions at the time of exposure. The initial concentration of target cells, that are the epithelial cells expressing the ACE2 receptor, is expressed as 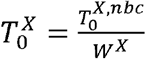 where 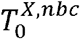 is the initial number of cells and *W*^*X*^ is the volume of distribution of the compartment of interest (see “Consideration of the volume of distribution”). Each animal was exposed to 1×10^6^ pfu of SARS-CoV-2 representing 2.19×10^10^ virions. Over the total inoculum injected (5 mL), 10% (0.5 mL) and 90% (4.5 mL) of virions were respectively injected by the intra-nasal route and the intra-tracheal route leading to the following initial concentrations of the incoculm within each compartment: 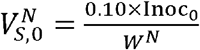 and 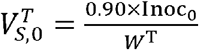, with Inoc_0_ the number virions injected via the inoculum.

Using the gRNA and sgRNA viral loads, we estimated the viral infectivity, the viral production rate and the loss rate of infected cells (**Table S2**). To account for inter-individual variability and covariates, each of those three parameters was described by a mixed-effect model and jointly estimated between the two compartments as follows:

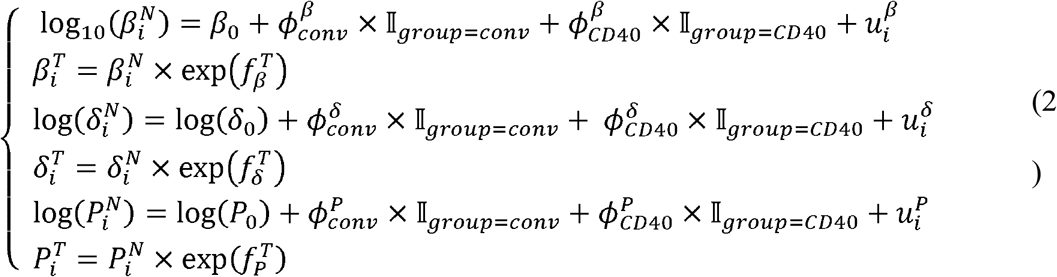

with 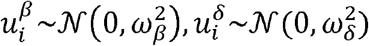 and 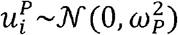, where *β*_0_,log(*δ*_0_) and log(*P*_0_) are the fixed effects, 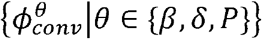 and 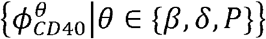 are respectively the regression coefficients related to the effects of the group of convalescent and αCD40.RBD vaccinated animals for the parameters β, δ and P, and 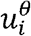 is the individual random effect for the parameter θ, which supposedly normally distributed with variance 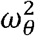.

In practice, after selection (see “Parameter estimation”), only random effects and group effects on the parameters β and δ were kept, fixing 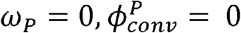 and 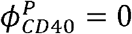. In addition, the estimation of several models identified the viral production rate P as the single parameter taken different values in nasopharynx and trachea 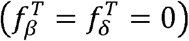. For the observation model, the log_10_-transformed genomic and subgenomic viral loads of the *i*th animal at the *j*th time point in the compartment X (nasopharynx or trachea), labelled 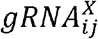 and 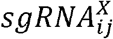 respectively, were described by the following equations:

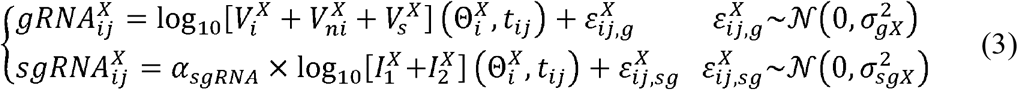

where 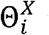 is the set of parameters of the subject *i* for the compartment X and ε are the additive normally distributed measurement errors.

### Consideration of the volume of distribution

To define the concentration of inoculum within each compartment after injection, nasopharyngeal and tracheal volumes of distribution, labelled *W*^*N*^ and *W*^*T*^ respectively, were requested. Given the estimated volumes of the trachea and the nasal cavities in four monkeys similar to our 18 macaques (**Figure S9A-C**) and the well documented relationship between the volume of respiratory tract and animal weights *(65)*, the volume of distribution of each compartment was defined as a step function of NHP weights:

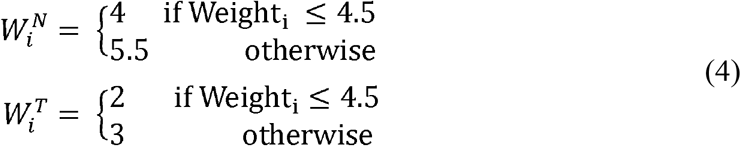

Where Weight_i_ is the weight of the monkey *i* in kgs. Using equation (4) and weights of our 18 NHPs (mean= 4.08; [Q1; Q3] = [3.26; 4.77]), we estimated W^T^ = 2 and W^N^ = 4mL for a third of them (n=12) (**Figure S9D**), leading to the initial concentration of target cells 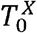 (see “Viral dynamics modeling” for equation) fixed at 3.13×10^4^ cells.mL^-1^ and 1.13×10^4^ cells.mL^-1^ in nasopharynx and trachea respectively. Similarly, their initial concentrations of challenge inoculum 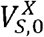 were fixed at 5.48×10^8^ copies.mL^-1^ and 9.86×10^9^ copies.mL^-1^ in nasopharynx and trachea resp. For the last third of NHPs (n=6), W^T^ = 3 and W^N^ = 5.5 mL leading to 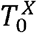 fixed at 2.27×10^4^ cells.mL^-1^ in nasopharynx and 7.50×10^3^ cells.mL^-1^ in trachea while 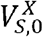 was fixed at 3.98×10^8^ copies.mL^-1^ in nasopharynx and 6.57×10^9^ copies.mL^-1^ in trachea. Through this modeling, we assumed a homogenous distribution of injected virions and target cells within nasopharyngeal and tracheal compartments. In addition, the natural downward flow of inoculum towards lungs, at the moment of injection, was indirectly taken into account by the parameter of inoculum clearance, c_i_.

### Parameter estimation

Among all parameters involved in the three layers of the mechanistic model, some of them have been fixed based on experimental settings and/or literature. That is the case of the proportion of infectious virus (μ) that has been fixed at 1/1000 according to previous work *(28)* and additional work (results not shown) evaluating the stability of the model estimation according to the value of this parameter. The initial number of target cells, that are the epithelial cells expressing the ACE2 receptor, 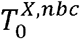 was fixed at 1.25×10^5^ cells in the nasopharynx and 2.25×10^4^ cells in trachea *(28)* (**Table S2**). The duration of the eclipse phase (*1/k*), the clearance of the inoculum (*c*_*i*_) and the clearance of the virus produced *de novo* (*c*) were estimated by profile likelihood. Although available data did not allow the direct estimation of these three parameters, the use profile likelihood enabled the exploration of various potential values for *k, c* and *c*_*i*_ In a first step, we explored the 18 models resulting from the combination of 3 values of *k* ∈ {1,3,6}day^-1^ and 6 values for *c*∈{1,5,10,15,20,30} day^-1^, assuming that the two parameters of virus clearance were equal, as first approximation. As shown in **Table S3**, an eclipse phase of 8 hours (k=3) and virus clearance higher than 15 virions per day led to lowest values of −2log-likelihood (−2LL, the lower the better). In a second step, we fixed the parameter *k* at 3 day^-1^ and estimated the 70 models resulting from the combination of 10 values for *c* ∈{1,2,3,4,5,10,15,20,25,30} day^-1^ and 7 values for *c*_*i*_ ∈ {1,5,10,15,20,25,30} day^-1^ (**Table S4**). The distinction of the two parameters of free virus clearance enabled to find much lower half-life of inoculum (∼50 minutes) than half-life of virus produced *de novo* (∼5.55 hours), with c=3 day^-1^ compared to c_i_=20 day^-1^.

Once all these parameters have been fixed, the estimation problem was restricted to the determination of the viral infectivity *β*, the viral production rate *P*, the loss rate of infected cells *δ* for each compartment, the parameter *α*_*vlsg*_ in the observation model, regression coefficients for groups of intervention (*ϕ*_conv_,*ϕ*_DC40_) and standard deviations for both random effects (*ω)* and error model (σ). The estimation was performed by Maximum likelihood estimation using a stochastic approximation EM algorithm implemented in the software Monolix (http://www.lixoft.com). Selection of the compartment effect on parameters (β, δ, P) as well as random effects and covariates on the statistical model (2) was performed by the estimation of several models that were successively compared according to the corrected Bayesian information criterion (BICc) (to be minimized). After the removal of random effect on the viral production (*ω*_*p*_= 0) allowing the reduction of the variance on the two other random effects, all combinations of compartment effects were evaluated, leading to the final selection of a single effect on *P* 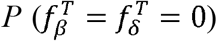. Then, the effect of group intervention was independently added on model parameters among *β, δ, P* and *c*. Once the group effect on the viral infectivity identified as the best one, the addition of a second effect on the remaining parameters was tested, resulting in the selection of the loss rate of infected cells. Finally, the irrelevance of the addition of a third effect was verified.

### Exchange of viruses between nasopharynx and trachea compartments

The possibility for viruses to migrate from nasopharyngeal to tracheal compartment and vice versa was tested. To this end, equations of infectious (V_i_) and non-infectious (V_ni_) viruses in equation (1) between the two compartments were linked as follows:

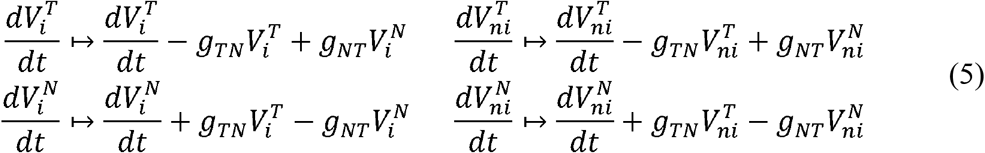

with the arrow symbolizing the modification of the equations defined in (1) and where g_TN_ and g_NT_ are the positive constant rates of exchange from trachea to nasopharynx and vice versa, respectively. Data described in the main article were too much sparse to estimate either bidirectional or at least one of the two unidirectional transfers defined by g_TN_ and g_NT_, additional data were used. Two naive macaques were exposed to the same dose (1×10^6^ pfu) of SARS-CoV-2 than our 18 monkeys but were inoculated via intra-gastric route (4.5mL) instead of intra-tracheal route. Similarly to our study, the viral gRNA dynamics in both tracheal and nasopharyngeal compartments were repeatedly measured during the 20 days following the challenge (**Figure S9E**). The model resulting from equation (5) was used to fit these data, considering all parameters as fixed (see **Table S2**), except for g_TN_ and g_NT._ The estimation of multiple models on those 2 animals tended to conclude that only an unidirectional transfer of viruses from the nasopharyngeal to the tracheal compartment should be explored, with an estimation of g_TN_ ranging from 0.9 to 2.5 day^-1^. However, the use of those fixed values in the estimation of the model on our 18 animals led irremediably to the degradation of the model with an increase of more than 2 points of BICc. An estimation of this parameter by profile likelihood (results not shown), resulting in a strictly decreasing profile of the likelihood (the higher the better), was not more conclusive. Consequently, we fixed the values of g_TN_ and g_NT_ at 0 day^-1^.

### Algorithm for automatic selection of biomarkers as CoP

After identifying the effect of the group of intervention on both the viral infectivity (β) and the loss rate of infected cells (δ), we aimed at determining whether some immunological markers quantified in the study could capture this effect. To this end, we developed a classic stepwise data-driven automatic covariate modelling method (**Figure S10**). The specificity of this method is the possibility to add either time-dependent or constant covariates in the model.

At the initialization step (k=0) (see **Figure S10**), the algorithm requests 3 inputs: (1) a set of potential *M* covariates, labelled Marker *m* ∈ {1, …,*M*} (e.g., immunological markers); (2) a set of *P* parameters on which covariates could be added, labeled *θ*_*p*_ for *p* ∈{1,…,*P*}(e.g. β and δ); and (3) an initial model (e.g., the model without covariates), labelled *M*^*0*^, with 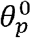 the definition of the parameter θ_*p*_. At each step k>0, we note *M* ^*k-1*^ the current model resulting in the model built in the step k-1. Then each combination of markers and parameters that have not already been added in *M* ^*k-1*^, labelled r (*r* ∈{Marker m ⊗*θ*^*p*^ ∉ *M*^*k*−1^ | *m* ∈ {1,… *M*},*p* ∈ {1,… *P*}}), are considered and tested in an univariate manner (each relation r is independently added in *M* ^*k-1*^ and ran). To this end, the parameter θ_*p*_ involved in this relationship r is modified as 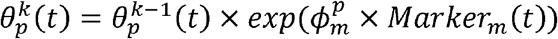, where 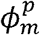 is the regression coefficient related the marker, while other parameters remain unchanged 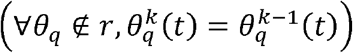. Once all these models evaluated, the one with the optimal value of a given criteria defining the quality of the fits (e.g., the lowest BICc value) is selected and compared to the model *M* ^*k-1*^. If its criteria value is better than the one found for *M* ^*k-1*^, then this model is defined as the new current model, *M* ^*k*^, and the algorithm moves to the step k+1. Otherwise, the algorithm stops. The algorithm can also be stopped at the end of a fixed number of step K.

The objective of this algorithm being to identify mechanistic correlates of protection, at each step, the selected model should respect, in addition to the best fits criteria, the 2 other criteria defining mCoP meaning the ability to capture the effect of the group of intervention and the ability to better explain the variability on individual parameters than the model adjusted on the group effect. To this end, we verify that in the selected model additionally adjusted on the group of intervention, the group effect appears as non-significantly different from 0 using a Wald-test. Then, we check that the variances of random effects in the selected model are well lower or equal to the ones obtained in the model adjusted only on the group effect.

### Quantification and statistical analysis

Statistical significance of the effect of groups in model estimation is indicated in the tables by stars: *, p < 0.05; **, p < 0.01; ***, p < 0.001 and were estimated by Wald test (Monolix^®^ software version 2019R1). In addition, statistical significance between viral loads in the two published studies (Brouwer et al, Cell 2021; Marlin et al., Nat Com 2021) in the control group were estimated by Welch two-sample t-test (R version 3.6.1) and are indicated in the supplementary file by p value. Model parameters were estimated with the SAEM algorithm (Monolix^®^ software version 2019R1).

Graphs were generated using R version 3.6.1 and Excel 2016 and details on the statistical analysis for the experiments can be found in the accompanying figure legends. Horizontal red dashed lines on graphs indicate assay limit of detection.

## Supplementary Materials

Fig. S1. Viral dynamics after the first exposure to SARS-CoV-2 and biomarker measurements from the first to the second exposure to SARS-CoV-2.

Fig. S2. Subgenomic viral dynamics after the second exposure to SARS-CoV-2.

Fig. S3. Modelling of the viral dynamics using mechanistic model.

Fig. S4. Antibody measurements after the second exposure to SARS-CoV-2.

Fig. S5. Antigen-specific T-cell responses in NHPs after the second exposure to SARS-CoV-2.

Fig. S6. Cytokines and chemokines in the plasma in NHPs after the second exposure to SARS-CoV-2.

Fig. S7. Immune markers selection and Basic reproduction number.

Fig. S8. The second study testing two-component spike nanoparticle vaccine. Fig. S9. Modelling of the dynamics of viral replication.

Fig. S10. Flow chart of the algorithm for automatic selection of covariate.

Table S1. Criteria defining RBD-ACE2 binding inhibition or neutralization measured on live cells with luciferase marker as mechanistic correlate of protection of the effect of the vaccine on new cell infection.

Table S2. Model parameters for viral dynamics in both the nasopharynx and the trachea estimated by the model adjusted for groups of intervention.

Table S3. Values of −2LL estimated on models with viral clearance (c=ci) and eclipse phase rate k fixed at different values.

Table S4. Values of −2LL estimated on models with inoculum clearance c_i_ and clearance of virus de novo produced c fixed at different values.

## Acknowledgments

We would like to thank J. Guedj and O. Terrier for fruitful discussions on the model definition. We thank S. Langlois, J. Demilly, N. Dhooge, P. Le Calvez, M. Potier, J. M. Robert, T. Prot, and C. Dodan for the NHP experiments; L. Bossevot, M. Leonec, L. Moenne-Loccoz, M. Calpin-Lebreau, and J. Morin for the RT-qPCR, ELISpot and Luminex assays, and for the preparation of reagents; A-S. Gallouët, M. Gomez-Pacheco and W. Gros for NHP T-cell assays and flow cytometry; B. Fert for her help with the CT scans; M. Barendji, J. Dinh and E. Guyon for the NHP sample processing; S. Keyser for the transports organization; F. Ducancel and Y. Gorin for their help with the logistics and safety management; I. Mangeot for here help with resources management and B. Targat contributed to data management. The monkey and syringe pictures in Fig.1 was created with BioRender.com. This work was supported by INSERM and the Investissements d’Avenir program, Vaccine Research Institute (VRI), managed by the ANR under reference ANR-10-LABX-77-01. MA has been funded by INRIA PhD grant. The Infectious Disease Models and Innovative Therapies (IDMIT) research infrastructure is supported by the “Programme Investissements d’Avenir”, managed by the ANR under reference ANR-11-INBS-0008. The Fondation Bettencourt Schueller and the Region Ile-de-France contributed to the implementation of IDMIT’s facilities and imaging technologies used to define volume of respiratory tract. The NHP study received financial support from REACTing, the Fondation pour la Recherche Medicale (FRM; AM-CoV-Path). We thank Lixoft SAS for their support. Numerical computations were in part carried out using the PlaFRIM experimental testbed, supported by Inria, CNRS (LABRI and IMB), Université de Bordeaux, Bordeaux INP and Conseil Régional d’Aquitaine (see https://www.plafrim.fr).

## Author contributions

Conceptualization: MA, RT, RLG, YL, RM

Methodology: MA, MP, RT

Software: MA, MP

Validation: MA, RT, MP

Investigation: RM, SC, NK, SC, TN, BD, MS, NDB, MC, PM, CL, AW

Resources: RM, SC, NK, SC, TN, BD, MS, NDB, MC, PM, CL, AW, OS, RWS, RLG, YL, MA

Writing – Original draft: RT, MA, YL, RLG, RM, MP

Writing – Review & Editing: All

Visualization: MA, RM, NK, TN, MP

Supervision: RT, RLG, YL, MP

Project administration: RT, RLG, YL

Funding acquisition: RT, RLG, YL

## Data and materials availability

No unique reagents were generated for this study.

Data from the studies ^30^ and ^37^ are available upon request. Data from the study ^24^ are available as supplementary material online.

The original Monolix code is available and free-of-cost on github (Inria SISTM Team) at the following link: https://github.com/sistm/SARSCoV2modelingNHP.

