## Supplemental Information for "SARS-CoV-2 mechanistic correlates of protection: insight from modelling response to vaccines"

Supplementary information for: SARS-CoV-2 mechanistic correlates of protection: insight from modelling response to vaccines

**Authors:** Marie Alexandre, Romain Marlin, Mélanie Prague, Séverin Coleon, Nidhal Kahlaoui, Sylvain Cardinaud, Thibaut Naninck, Benoit Delache, Mathieu Surenaud, Mathilde Galhaut, Nathalie Dereuddre-Bosquet, Mariangela Cavarelli, Pauline Maisonnasse, Mireille Centlivre, Christine Lacabaratz, Aurelie Wiedemann, Sandra Zurawski, Gerard Zurawski, Olivier Schwartz, Rogier W Sanders, Roger Le Grand, Yves Levy, Rodolphe Thiébaut

*Corresponding author:	Prof Rodolphe Thiébaut

Bordeaux University, Departement of Public Health

146 Rue Leo Saignat, 33076 Bordeaux Cedex, France

**Supplementary Figures and Tables**

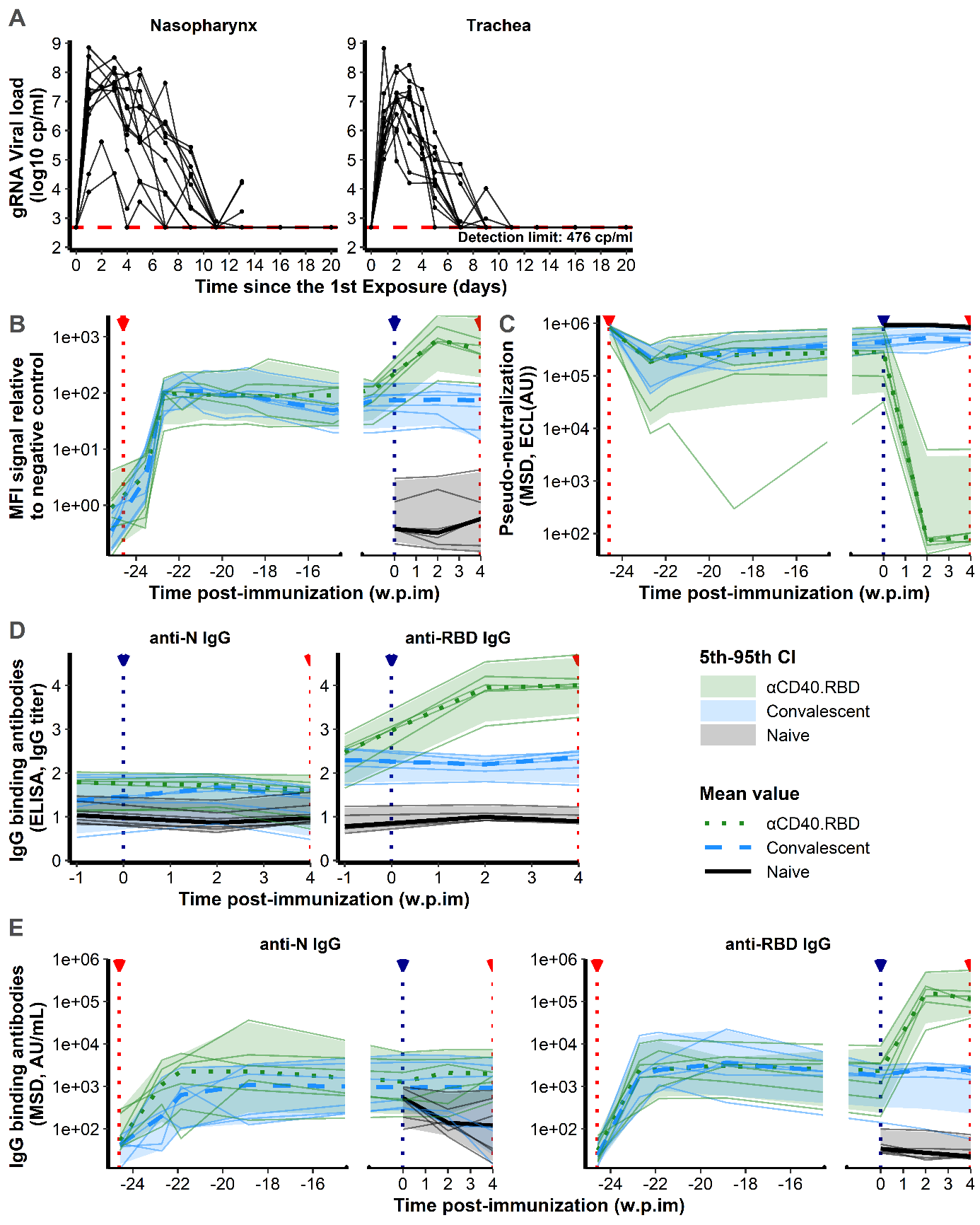

**Fig. S1. Viral dynamics after the first exposure to SARS-CoV-2 and biomarker measurements from the first to the second exposure to SARS-CoV-2.**

**(A)** Individual log10 transformed gRNA viral load dynamics in nasopharyngeal (left) and tracheal (right) swabs after the initial exposure to SARS-CoV-2 in naive macaques (n=12). Solid lines represent individual values. Horizontal red dashed lines indicate the limit of quantification. **(B)** Relative MFI of IgG binding to SARS-CoV-2 Spike protein, measured using a Luminex-based serology assay, in serum samples, after the initial exposure to SARS-CoV-2. **(C)** Quantification of antibodies inhibiting the attachment of Spike protein to ACE2 receptor in NHP serum, measured by the Mesoscale Discovery (MSD, Rockville, MD) pseudo-neutralization assay. Results are expressed as ECL (ECL, Electro-chemioluminescence) in AU. **(D)** Quantification of SARS-CoV-2 IgG binding N or RBD domain measured in the serum of NHPs titrated by ELISA assay. Results are expressed in IgG titer. **(E)** Quantification of SARS-CoV-2 IgG binding N and RBD domain measured in the serum of NHPs using a multiplexed solid-phase chemiluminescence assay. Results are expressed in AU/mL. (**B-E)** Results are obtained after the initial exposure to SARS-CoV-2 at -24.9 weeks post-immunization (w.p.im) in convalescent (n=6, blue, dashed line) and αCD40.RBD-vaccinated convalescent (n=6, green, dotted line) animals and at 4 w.p.im in naive (n=6, black, solid line) animals. Thin lines represent individual values. Thick lines indicate medians within each group and shaded areas indicate 5^th^-95^th^ confidence intervals. The red (-24.6 and 4.0 w.p.im) and blue (0 w.p.im) lines highlight viral exposure and vaccination respectively.

**Fig. S2. Subgenomic viral dynamics after the second exposure to SARS-CoV-2**

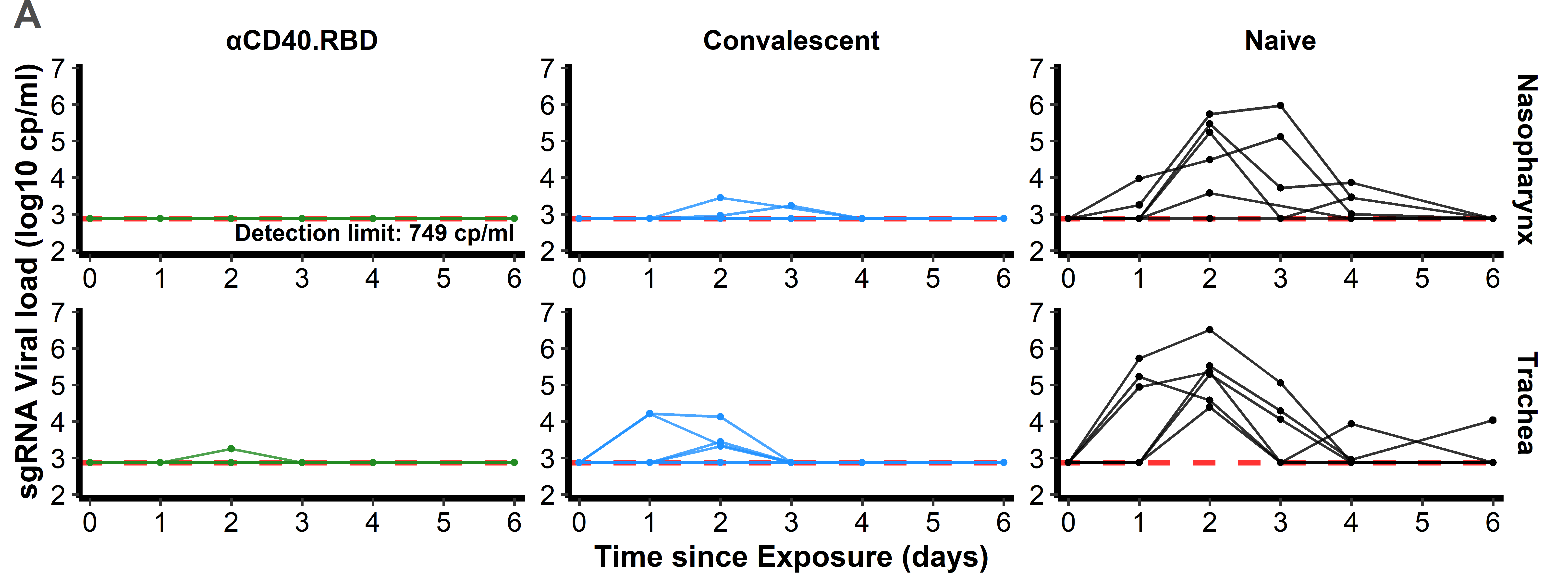

Individual log10 transformed subgenomic (gRNA) viral load dynamics in nasopharyngeal (top) and tracheal (bottom) swabs after the initial exposure to SARS-CoV-2 in naive macaques (n=6, black, right) and after the second exposure in convalescent (n=6, blue, middle) and αCD40.RBD-vaccinated convalescent (n=6, green, left) groups. Horizontal red dashed lines indicate the limit of quantification.

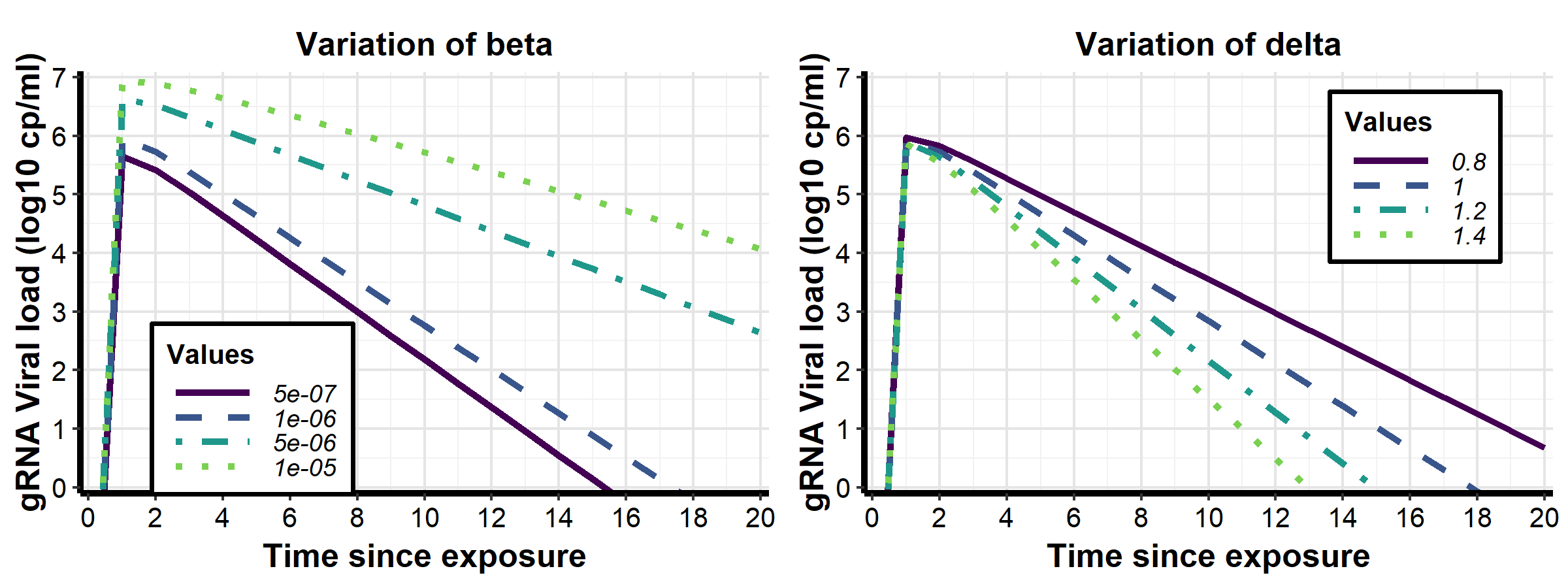

**Fig. S3. Modelling of the viral dynamics using mechanistic model**

Examples simulated genomic viral load dynamics for different values of viral infectivity (β, left) or loss rate of infected cells (δ, right) showing the effect of either blocking de novo infection or promoting the destruction of infected cells on viral dynamics profile. Except for β or δ, all other parameters were fixed at a given value.

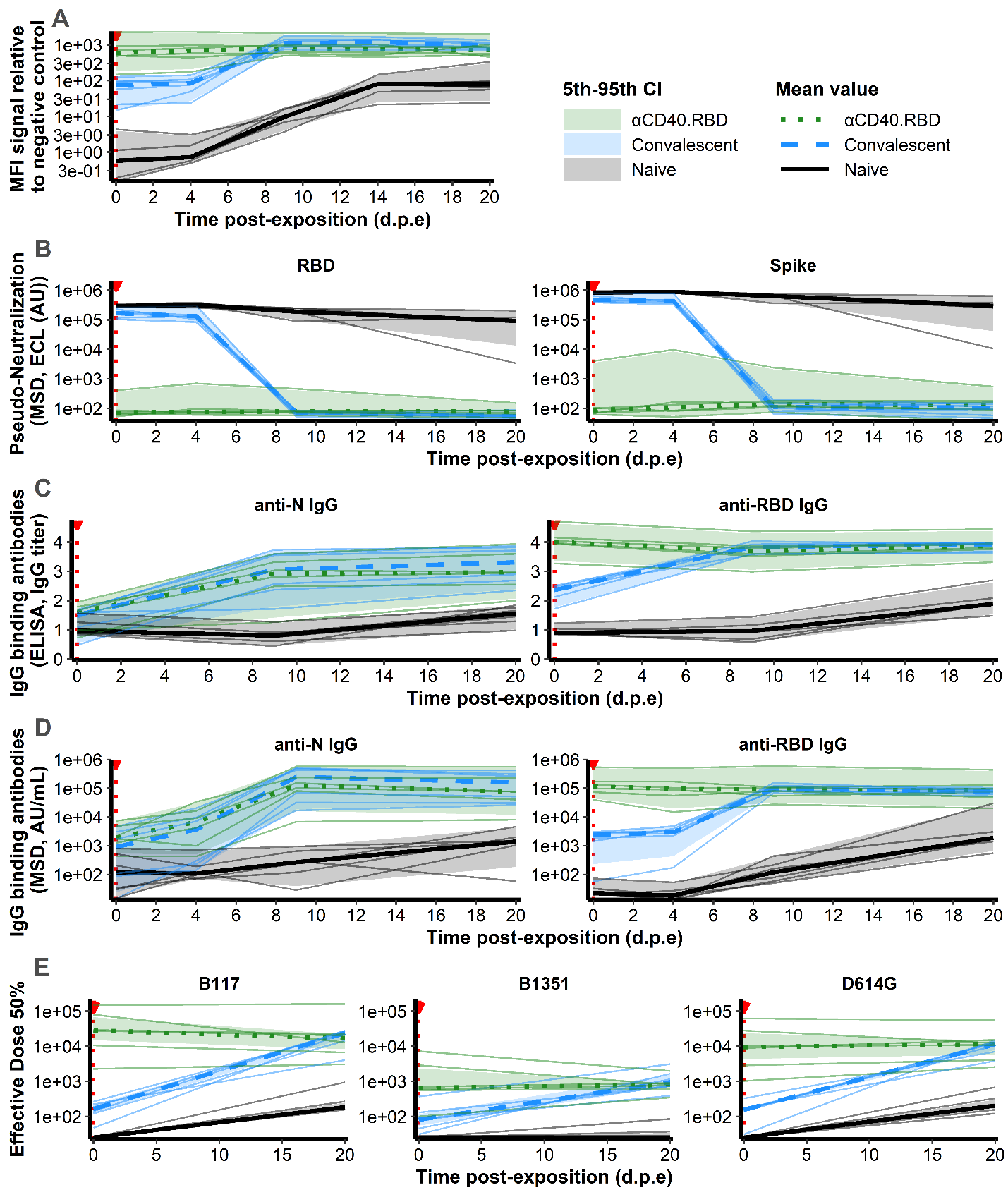

**Fig. S4. Antibody measurements after the second exposure to SARS-CoV-2**

**(A)** Relative MFI of IgG binding to SARS-CoV-2 Spike protein, measured using a Luminex-based serology assay, in serum samples, after the second exposure to SARS-CoV-2. **(B)** Quantification of antibodies inhibiting that attachment of RBD domain or Spike protein to ACE2 receptor in NHP serum, measured by the Mesoscale Discovery (MSD, Rockville, MD) pseudo-neutralization assay, after the second exposure to SARS-CoV-2. Results are expressed as ECL, in AU. **(C)** Quantification of SARS-CoV-2 IgG binding N or RBD domain measured in the serum of NHPs titrated by ELISA assay, after the second exposure to SARS-CoV-2. Results are expressed in Ig titer. **(D)** Quantification of SARS-CoV-2 IgG binding N or RBD domain measured in the serum of NHPs using a multiplexed solid-phase chemiluminescence assay, after the second exposure to SARS-CoV-2. Results are expressed in AU/mL. **(E)** Quantification of neutralizing antibodies against B.1.1.7, B.1.351 and D614G SARS-CoV-2 strains measured in the serum of NHPs using S-Fuse neutralization assay, after the second exposure to SARS-CoV-2 (measured only at the exposure and 20 days post-exposure (d.p.e)). Results are expressed as ED50 (Effective dose 50%). **(A-E)** Results are obtained after the initial exposure to SARS-CoV-2 in naive macaques (n=6, black, solid line) and after the second exposure in convalescent (n=6, blue, dashed line) and αCD40.RBD-vaccinated convalescent (n=6, green, dotted line) animals. Thin lines represent individual values. Thick lines indicate medians within each group and shaded areas indicate 5th-95th confidence intervals. Red dotted vertical lines highlight the viral exposure.

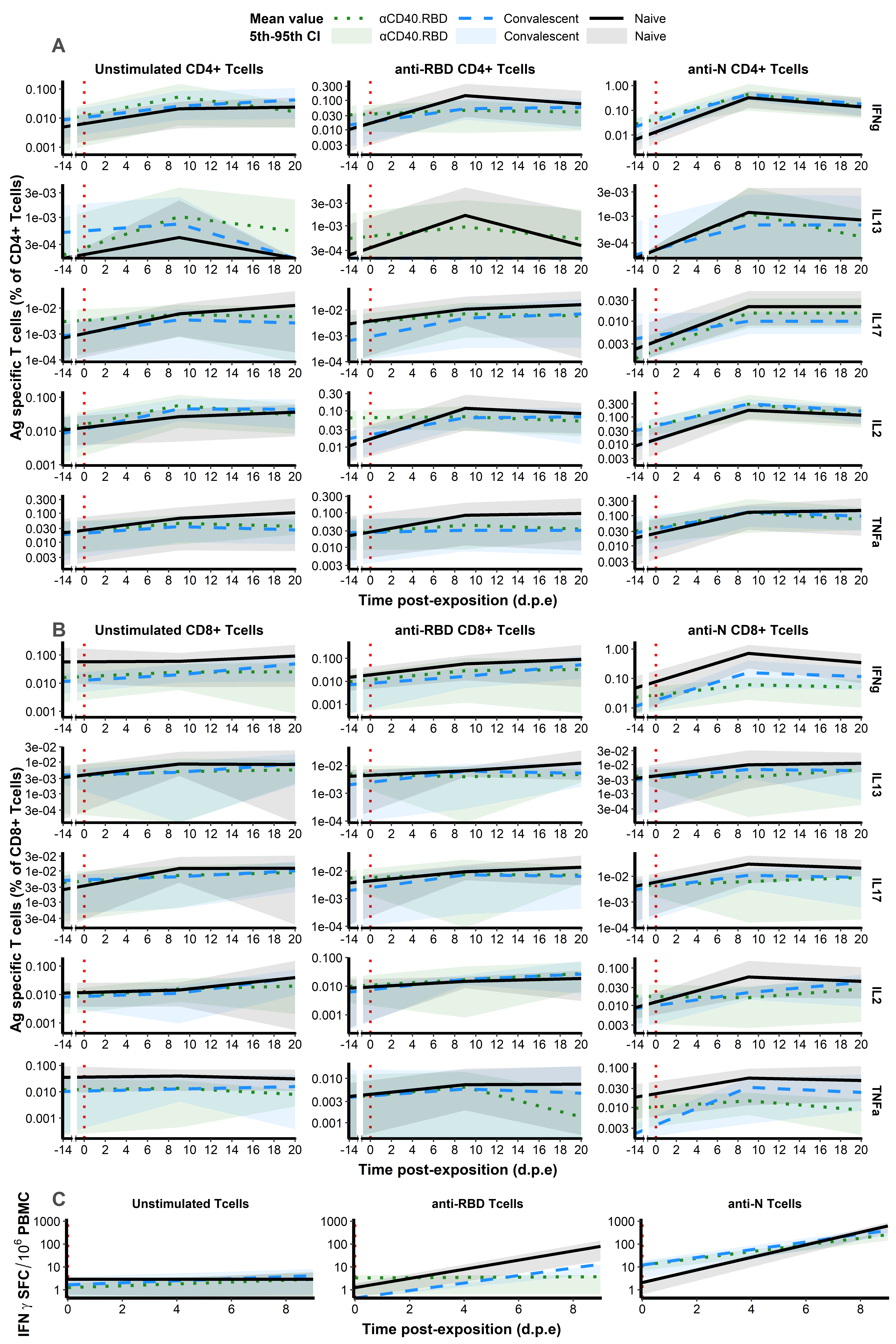
**Fig. S5. Antigen-specific T-cell responses in NHPs after the second exposure to SARS-CoV-2**

**(A-B)** Frequency of IFNγ^+^ (fist line), IL-13^+^ (second line), IL-17^+^ (third line), IL-2^+^ (fourth line) or TNFα^+^ (fifth line) antigen-specific CD4^+^ Tcells (CD154^+^) and CD8^+^ Tcells (CD137^+^) in the total CD4^+^ Tcell (A) or CD8^+^ Tcell (B) population in NHP serum. PBMCs were stimulated *ex-vivo* overnight with medium (left), SARS-CoV-2 RBD (middle) or N (right) overlapping peptide pools. T-cell responses being not measures at the challenge, measured obtained 14 days pre-exposure were added. **(C)** Antigen-specific T-cell responses in NHPs. T-cells were analyzed by ELISpot after *ex-vivo* stimulation with SARS-CoV-2 RBD or N overlapping peptide pools and plotted as spot-forming cells (SFC) per 1.0x10^6^ PBMCSs. **(A-C)** Results are obtained after the initial exposure to SARS-CoV-2 in naive macaques (n=6, black, solid line) and after the second exposure in convalescent (n=6, blue, dashed line) and αCD40.RBD-vaccinated convalescent (n=6, green, dotted line) animals. Thin lines represent individual values. Thick lines indicate medians within each group and shaded areas indicate 5th-95th confidence intervals. Red dotted vertical lines highlight the viral exposure.

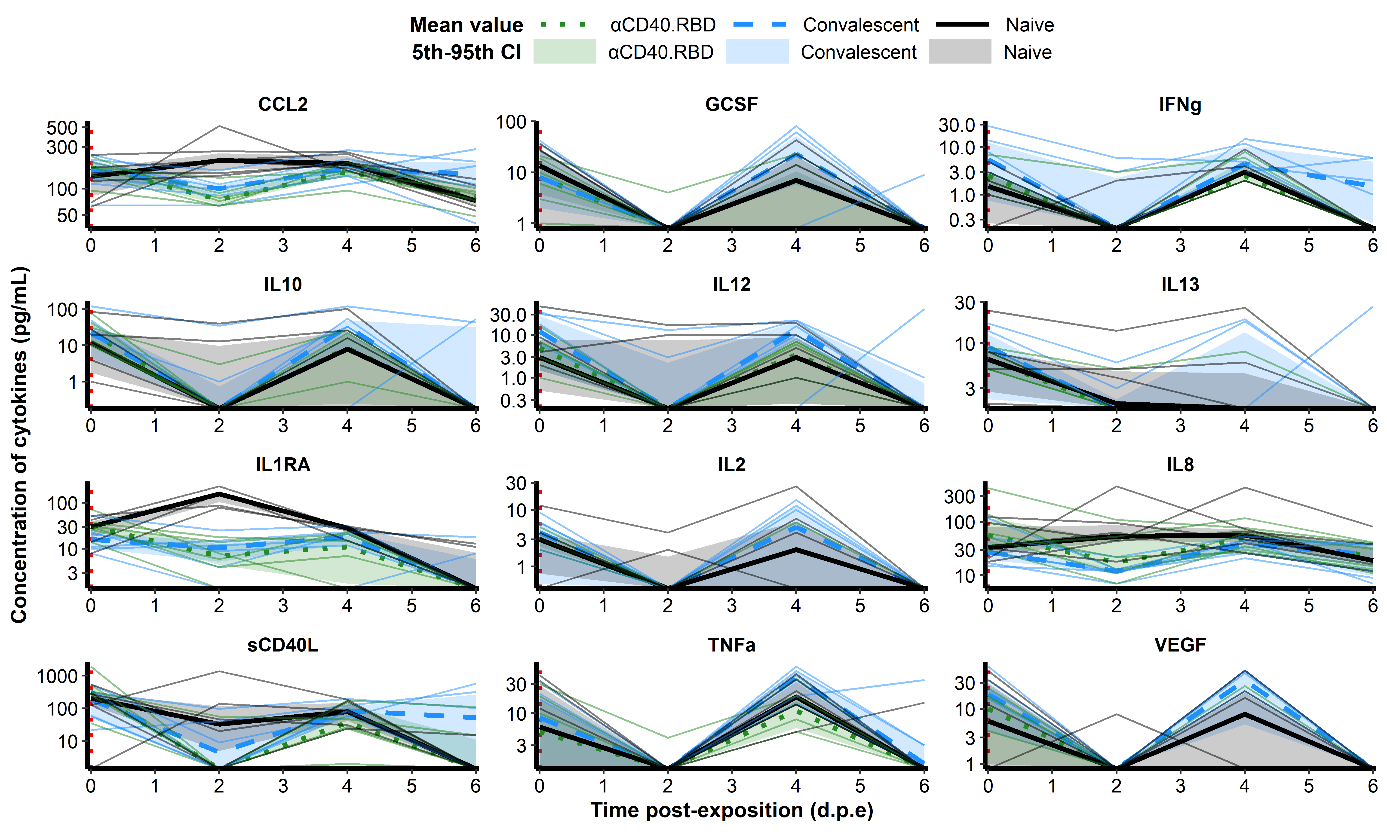

**Fig. S6. Cytokines and chemokines in the plasma in NHPs after the second exposure to SARS-CoV-2**

Plasma concentration of 12 cytokines and chemokines in pg/mL. Results are obtained after the initial exposure to SARS-CoV-2 in naive macaques (n=6, black, solid line) and after the second exposure in convalescent (n=6, blue, dashed line) and αCD40.RBD-vaccinated convalescent (n=6, green, dotted line) animals. Thin lines represent individual values. Thick lines indicate medians within each group and shaded areas indicate 5th-95th confidence intervals. Red dotted vertical lines highlight the viral exposure.

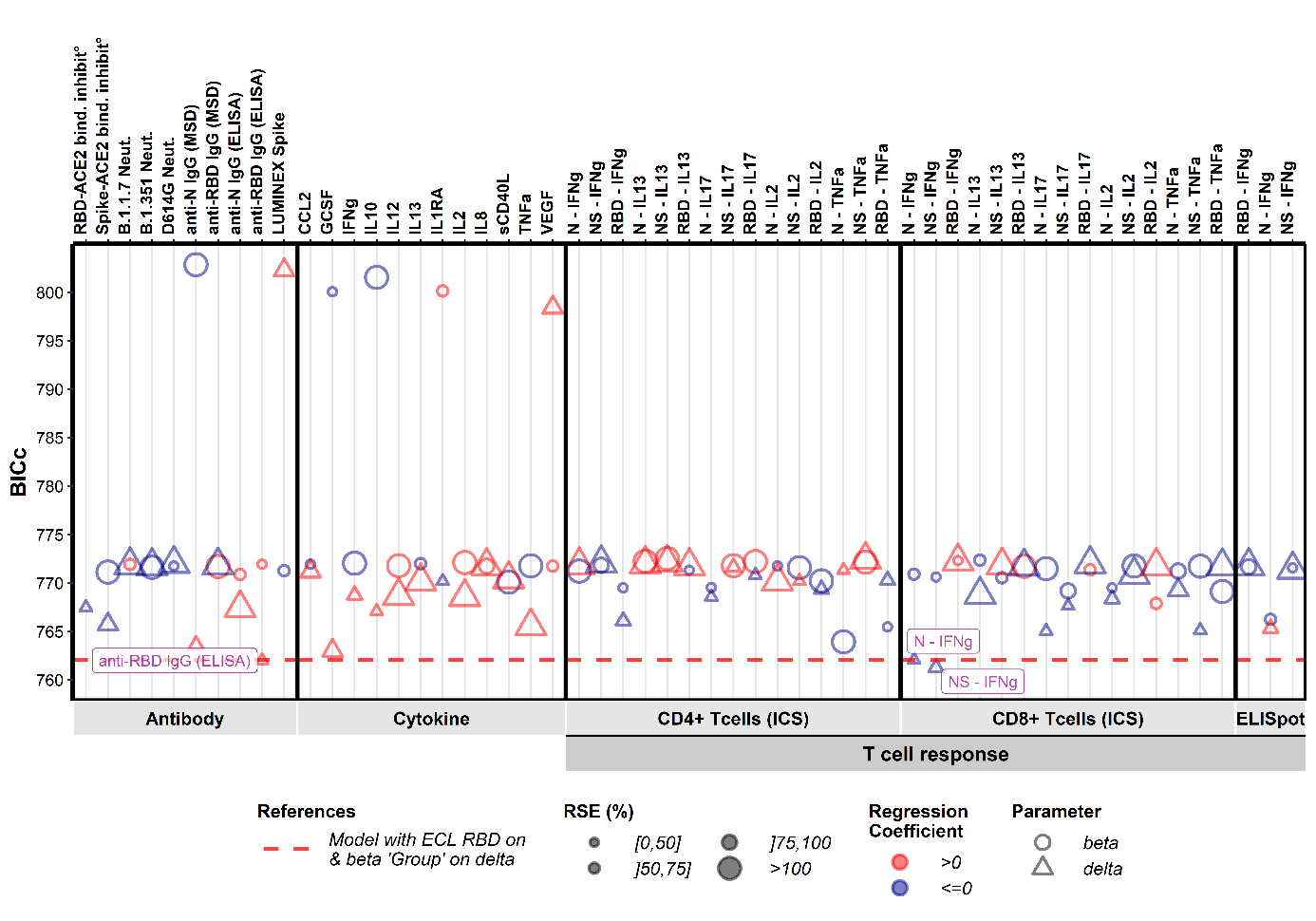

**A**

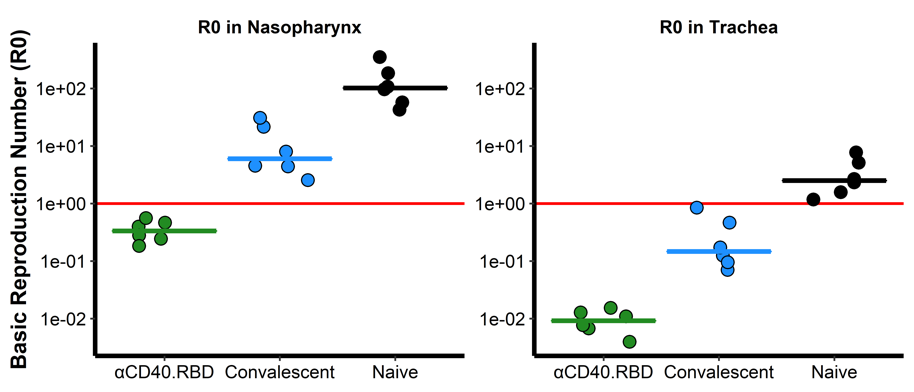

**B**

**Fig. S7. Immune markers selection and Basic reproduction number**

**(A)** *Systematic screening of effect of the markers (Step 2).* For every single marker, a model, already adjusted on viral infectivity with antibodies inhibiting the attachment of RBD domain to ACE2 receptor, has been fitted to explore whether it explains the variation of the parameter of interest better or as well than the model of reference. Parameters of interest were β, the infection rate of ACE2+ target cells and δ, the loss rate of infected cells. Models were compared according to the Bayesian Information Criterion (BIC), the lower being the better. The red dashed line represents the reference model that includes the group effect (naive/ convalescent/vaccinated) on the parameter δ and with adjustment of pseudo-neutralization on β. **(B)** *Reproduction rate at the time of exposure.* Model predictions of the reproduction rate at the time of exposure (R_0_) in the tracheal (right) and nasopharyngeal (left) compartments for naive (black), convalescent (blue) and αCD40.RBD-vaccinated convalescent (green) animals. The reproduction rate is representing the number of infected cells from one infected cell if target cells are unlimited. When this effective reproduction rate is below 1, it means that the infection is going to be cured. The values of R_0_ were estimated by the model with viral infectivity (β) and loss rate of infected cells (δ) adjusted on pseudo-neutralization and anti-RBD IgG binding antibodies titrated by ELISA assay respectively measured only at the time of challenge. Horizontal solid red lines highlight the threshold of the reproduction rate equals to one.

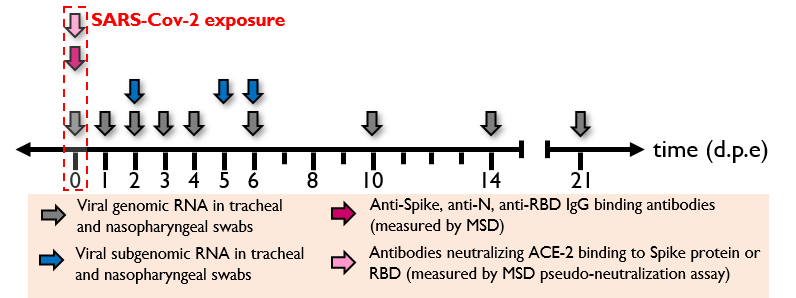

**B**

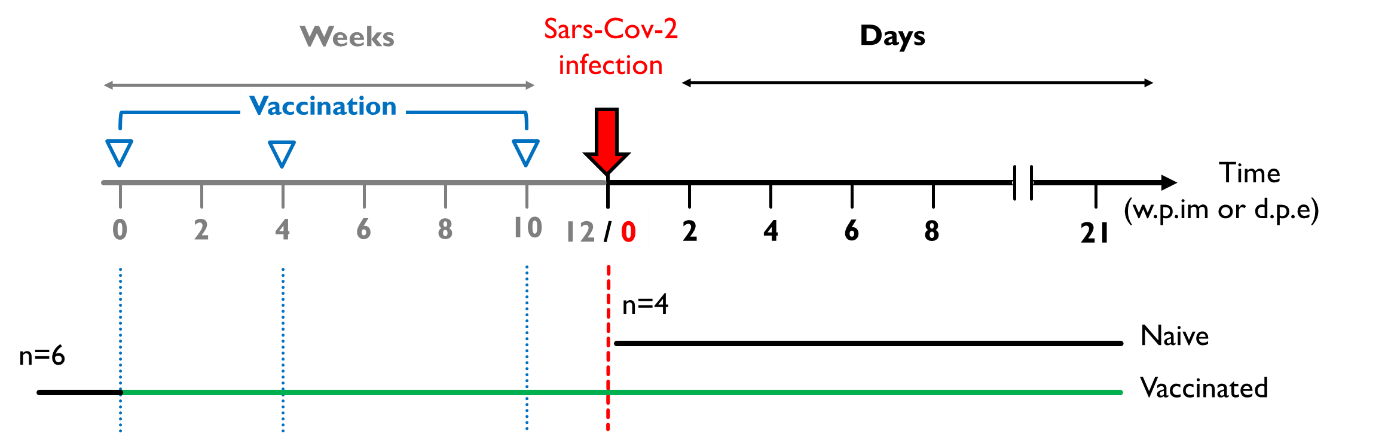

**A**

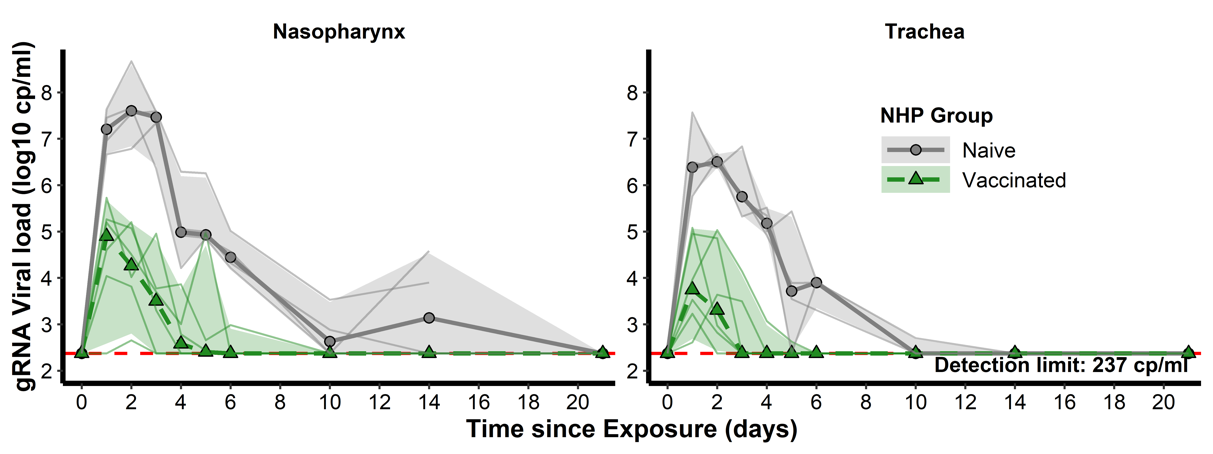

**C**

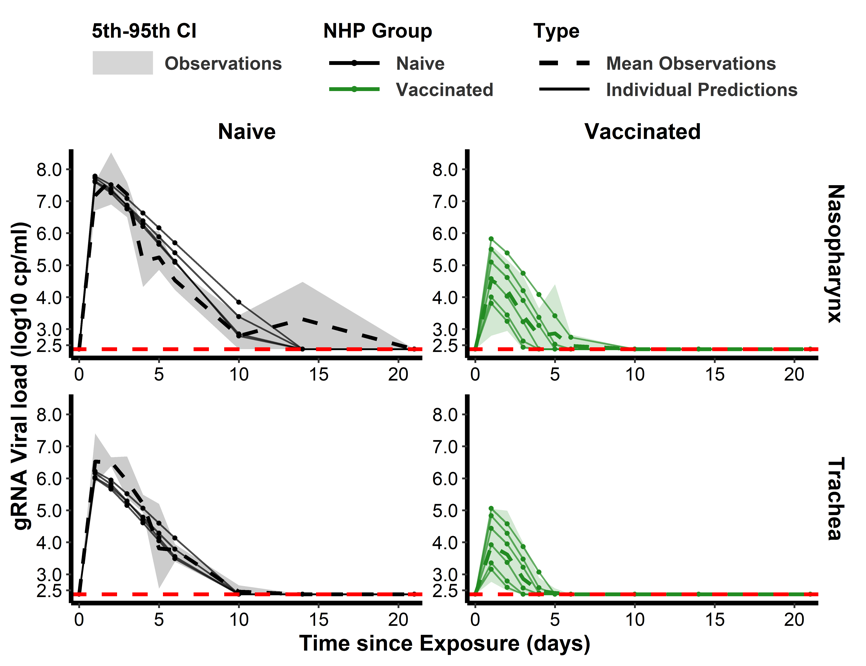

**D**

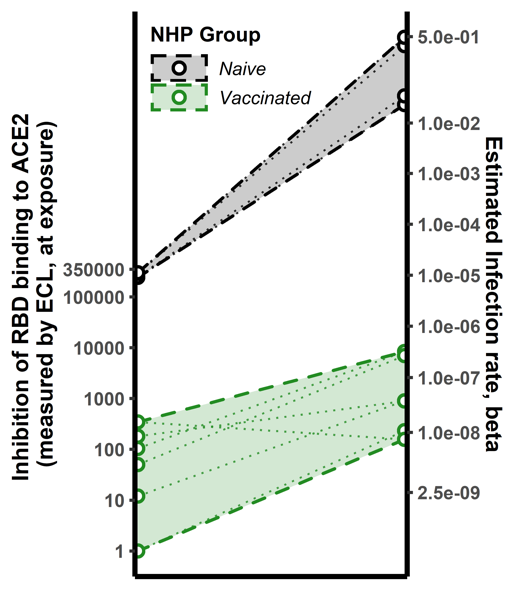

**E**

**Fig. S8. The second study testing two-component spike nanoparticle vaccine.**

(A) *Study design*. Cynomolgus macaques were randomly assigned in two experimental groups. Twelve, eight and two weeks post-infection with SARS-CoV-2 virus, six of them were successively immunized with 50 µg of SARS-CoV-2 S-I53-50NP vaccine. The four other animals received no vaccination. Two weeks after the final immunization, all monkeys were exposed to a total dose of 10^6^ pfu of SARS-CoV-2 virus via intra-nasal and intra-tracheal routes. (B) *Harvest times and measurements*. Nasopharyngeal and tracheal fluids were collected at 0, 1, 2, 3, 4, 5, 6, 10, 14 and 21 d.p.e while blood was taken at 0, 2, 4, 6, 10, 14 and 21 d.p.e. Genomic and subgenomic viral loads were measured by RT-qPCR. Anti-Spike, anti-RBD and anti-Nucleocapside (N) IgG were titrated using a multiplexed immunoassay developed by Mesoscale Discovery (MSD, Rockville, MD) and expressed in AU/mL. The MSD pseudo-neutralization assay was used to quantify antibodies neutralizing the binding of the spike protein and RBD domain to the ACE2 receptor and results were expressed in ECL. (C) Genomic viral load dynamics in nasopharyngeal and tracheal swabs after the exposure to SARS-Cov-2 in naive (black, solid line) and vaccinated (green, dashed line) animals. Thin lines represent individual values. Thick lines indicate medians within each group. (D) Model fit to the log10-transformed observed gRNA viral load in nasopharynx and trachea after the exposure to SARS-CoV-2 in naïve and vaccinated macaques. Solid thin lines indicate individual dynamics predicted by the model adjusted for groups. Thick dashed lines indicate mean viral load over time. (E) *Thresholds of inhibition of RBD-ACE2 binding.* Estimated infection rate of target cells ((copies/mL)^-1^day^-1^) according to the quantification of antibodies inhibiting RBD-ACE2 binding (ECL) at exposure for naive (black) and vaccinated (green) animals. Thin dotted lines and circles represent individual infection rates (right axis) and neutralizing antibodies (left axis). Thick dashed lines and dashed areas delimit the pseudo-neutralization / viral infectivity relationships within each group. (C,D) Horizontal red dashed lines represent the limit of quantification and shaded areas the 95% confidence intervals.

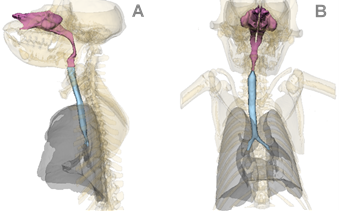

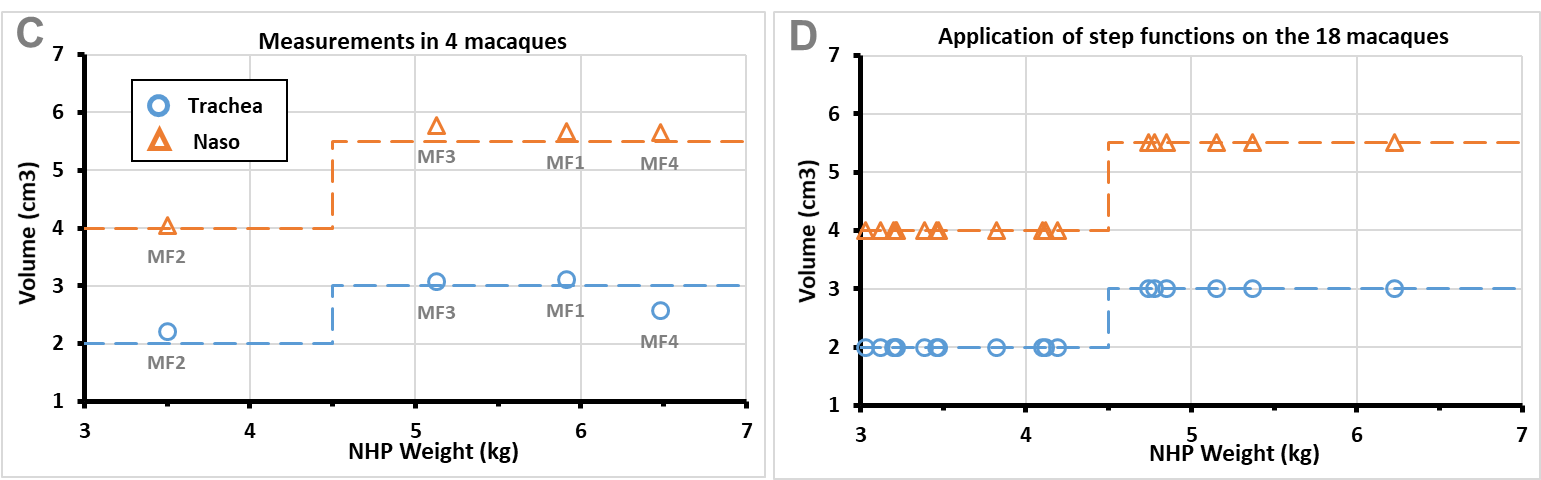

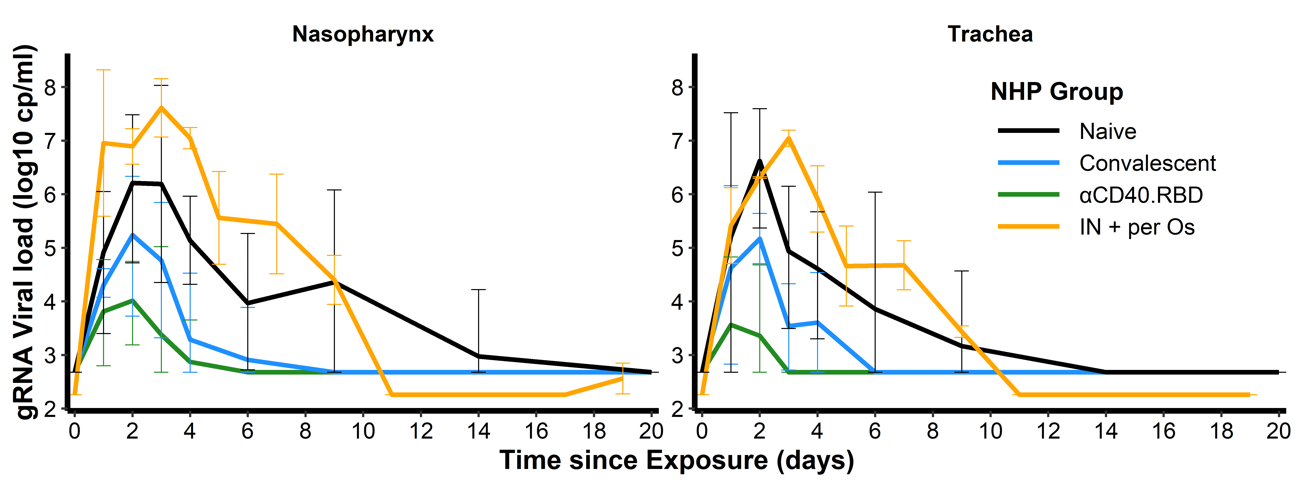

**E**

**Fig. S9. Modelling of the dynamics of viral replication**

**(A)** Sagittal view of the 3D representation of the NHP respiratory system. **(B)** Coronal view of the 3D representation of the NHP respiratory system. (A-B) Lungs are colored in grey, Trachea and Nasal regions in blue and purple respectively. **(C)** Relationship between the weights (in kgs) measured in 4 NHPs and the estimation of the volume of their tracheal (blue circles) and nasal (orange triangles) regions (in cm^3^). Measurements were obtained on NHPs similar to the 18 macaques of our study. Orange and blue dashed lines represent the step function used to describe this relationship with a breakpoint at 4.5 kg. **(D)** Volumes of the tracheal (blue circles) and nasal (orange triangles) regions estimated for the 18 macaques using the step function defined in the subfigure C and their weights. **(E)** Mean gRNA load dynamics in nasopharyngeal (left) and tracheal (right) swabs after the initial exposure to SARS-CoV-2 in naive macaques (n=6, black) and after the second exposure in convalescent (n=6, blue) and αCD40.RBD-vaccinated convalescent (n=6, green) macaques. Two additional macaques (IN + per Os, orange) were initially exposed to SARS-CoV-2 via intra-nasal (0.5mL of inoculum) and intra-gastric (4.5 mL) routes instead of intra-nasal (0.5 mL of inoculum) and intra-tracheal (4.5 mL) routes as defined in the study. Solid lines represent mean values and error bars indicate the 5^th^-95^th^ confidence intervals.

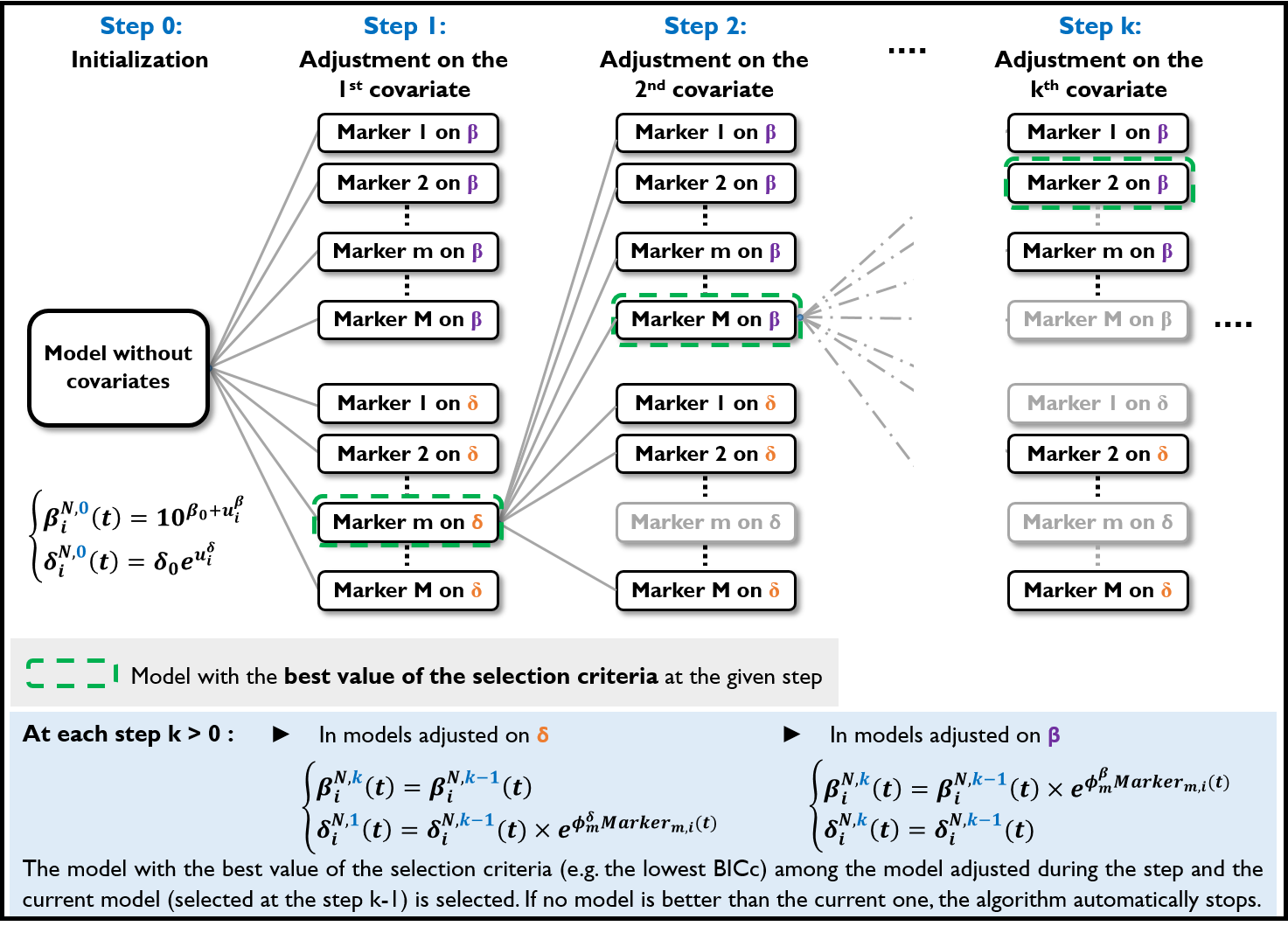

**Fig. S10. Flow chart of the algorithm for automatic selection of covariate.**

At the initialization step of our study, the model without covariates is considered as initial the model, all immunological markers are seen as potential covariates (Marker) and {β, δ} is defined as the set of parameters on which covariates can be added. At each, all the marker-parameter relationships that have not already been added to the current model are added in an univariate manner to this model and ran. Among all the tested models, the one with the optimal value of selection criteria (e.g. lowest BICc) is selected (green dashed rectangle) and compared to the current model. If this one is better, it becomes the new current model and the algorithm moves to the step k+1. Otherwise, the algorithm stops.

**Table S1.** Criteria defining RBD-ACE2 binding inhibition (studies testing two-component spike nanoparticle vaccine) or neutralization measured on live cells with luciferase marker (mRNA-1273) as mechanistic correlate of protection of the effect of the vaccine on new cell infection. Studies are labelled Study 1 and Study 2 for the main and the additional studies testing two-component spike nanoparticle vaccine, respectively and mRNA-1273 for the third study evaluating several doses of mRNA-1273 vaccine.

| **Model** | | **Study 1** | **Study 1** | **Study 2** | **mRNA-1273** |
| --- | --- | --- | --- | --- | --- |
|  |  | RBD-ACE2 binding inhibition on β | RBD-ACE2 binding inhibition on β | RBD-ACE2 binding inhibition on β | Neutralization on β |
| **Type of covariates added in the model** | | **Time-varying** | **Baseline** | **Baseline** | **Baseline** |
| **Criterion 1** | Without adjustment | 793.71 | 793.71 | 427.63 | 469.13 |
| Best fits (BICc Value) | Adjusted for Group | 772.34 | 772.34 | 404.80 | 468.70 |
|  | Adjusted for marker | 765.76 | 765.22 | 413.35 | 467.21 |
| **Criterion 2** | Adjusted for Group | **Conv*:*** -0.747** (0.255) | **Conv:** -0.747** (0.255) | **Vacc:** -6.01**** (0.8) | **10µg:** -0.267 (0.578) |
|  |  | **Conv-CD40:** -2.38*** (0.265) | **Conv-CD40:** -2.38*** (0.265) |  | **100µg:** -1.6** (0.621) |
| Group effect (Value (Sd)) | Adjusted for Group & marker | **Conv*:*** *0.00428 (1.71)* | **Conv:** 0.303 (0.864) | **Vacc:** 2.31 (1.18) | **10µg:** 0.105(0.646) |
|  |  | **Conv-CD40:** 0.1 (2.79) | **Conv-CD40:** 0.362 (1.14) |  | **100µg:** 3.10-4 (0.84) |
| **Criterion 3** | Adjusted for Group | **β:** 65.5 % | **β:** 65.5 % | **β:** 64.9 % | **β:** 18.5 % |
| Expl. variability for β or δ (relative %^1^) |  | **δ:** 54 % | **δ:** 54 % | **δ:** 58.2 % | **δ:** 4.6% |
|  | Adjusted for marker | **β:** 87.4 % | **β:** 82.7 % | **β:** 70.8 % | **β:** 15.4 % |
|  |  | **δ:** 27.1 % | **δ:** 31.6% | **δ:** -17% | **δ:** 19.1 % |
| *^1^ Relative percentage of inter-individual variability compared to variability obtained on model without any adjustment ; 100-(x*100)/ref* | | | | | |
| ** P < 0.05, **P < 0.01, ***P < 0.001, ****P<1e-4, Wald test for fold change different from 1* | | | | | |

**Table S2.** Model parameters for viral dynamics in both the nasopharynx and the trachea estimated by the model adjusted for groups of intervention.

| **Param.** | **Meaning** | **Value [95% CI]** | **Unit** |
| --- | --- | --- | --- |
| β | Infection rate in the naive group | 0.95 [0.18 ; 4.94] (x10^-6^) | (copies/ml)^-1^ day^-1^ |
|  | Fold change in the convalescent group | 0.18 [0.04 ; 0.88]** |  |
|  | Fold change in the Conv-CD40 group | 0.004 [0.001 ; 0.029]*** |  |
| δ | Loss rate of infected cells in the naive group | 1.04 [0.79 ; 1.37] | day^-1^ |
|  | Fold change in the convalescent group | 1.79 [1.21 ; 2.66]** |  |
|  | Fold change in the Conv-CD40 group | 1.80 [1.17 ; 2.75]** |  |
| P^N^ | Viral production rate in the naso. | 12.1 [3.15 ; 46.5] (x10^3^) | virions.(cell.day)^-1^ |
| P^T^ | Viral production rate in the trachea | 0.92 [0.39 ; 2.13] (x10^3^) | virions.(cell.day)^-1^ |
| α_vlsg_ | Infected cells and sgRNA viral load ratio | 1.39 [1.01 ; 1.76] | Virions.cell^-1^ |
| k | Eclipse rate | 3 | day^-1^ |
| c | Clearance of *de novo* produced viruses | 3 | day^-1^ |
| c_I_ | Clearance of inoculum | 20 | day^-1^ |
| µ | Percentage of infectious viruses | 10^-3^ |  |
| $T_{0}^{X,nbc}$ | Initial number of target cells | 1.25x10^5^ (Naso.)  2.25x10^4^ (Trachea) | cells |
| $\mathrm{Inoc}_{0}$ | Number of virions inoculated | 2.19x10^10^ | virions |
| ω_β_ | SD of random effect on log_10_ β | 0.366 [0.160 ; 0.571] |  |
| ω_δ_ | SD of random effect on δ | 0.170 [-0.089 ; 0.429] |  |
| σ_VLn_ | SD of error model gRNA in naso. | 1.27 [1.01 ; 1.53] |  |
| σ_VLt_ | SD of error model gRNA in trachea | 1.09 [0.90 ; 1.28] |  |
| σ_sgVLn_ | SD of error model sgRNA in naso | 1.41 [0.97 ; 1.85] |  |
| σ_sgVLt_ | SD of error model sgRNA in trachea | 1.62 [1.11 ; 2.13] |  |
| **Conv-CD40 group**: group of convalescent NHPs being vaccinated ; **SD**: Standard deviation ; ** P < 0.05, **P < 0.01, ***P < 0.001, wald test for fold change different from 1* | | | |

**Table S3.** Values of -2LL estimated on models with viral clearance (c=cI) and eclipse phase rate k fixed at different values.

| **c k** | **1** | **3** | **6** |
| --- | --- | --- | --- |
| **1** | 1285.29 | 1286.42 | 1289.54 |
| **5** | 871.38 | 864.50 | 866.80 |
| **10** | 773.74 | 764.29 | 768.18 |
| **15** | 749.67 | **738.71** | 742.12 |
| **20** | 749.44 | **738.40** | 740.98 |
| **30** | 750.00 | **739.51** | 741.34 |

**Table S4.** Values of -2LL estimated on models with inoculum clearance cI and clearance of virus de novo produced c fixed at different values. The eclipse phase rate was fixed at k=3 day^-1^.

| **c c_I_** | **1** | **5** | **10** | **15** | **20** | **25** | **30** |
| --- | --- | --- | --- | --- | --- | --- | --- |
| **1** | 1286.42 | 873.30 | 777.21 | 753.72 | 754.14 | 754.32 | 754.75 |
| **2** | 1286.70 | 864.90 | 760.94 | 734.03 | 734.14 | 733.94 | 734.95 |
| **3** | 1286.41 | 864.58 | 760.69 | 734.71 | **733.85** | 734.87 | 734.48 |
| **4** | 1286.33 | 864.22 | 761.78 | 735.91 | 735.16 | 735.42 | 736.14 |
| **5** | 1286.38 | 864.50 | 762.69 | 737.13 | 735.85 | 736.37 | 736.69 |
| **10** | 1286.55 | 865.48 | 764.29 | 738.38 | 737.56 | 737.89 | 738.13 |
| **15** | 1286.23 | 865.12 | 764.79 | 738.71 | 737.54 | 738.17 | 738.75 |
| **20** | 1285.96 | 865.19 | 764.78 | 738.86 | 738.40 | 738.25 | 739.30 |
| **25** | 1286.28 | 864.93 | 764.97 | 739.20 | 738.05 | 738.37 | 739.34 |
| **30** | 1286.45 | 864.77 | 765.16 | 739.01 | 738.13 | 738.58 | 739.51 |
